# Unveiling novel mechanisms of strobilurin resistance in the cacao pathogen *Moniliophthora perniciosa*

**DOI:** 10.1101/2024.11.14.623591

**Authors:** P.F.V. Prado, C.V.C. Mendes, B.A. Pires, G.L. Fiorin, P. Mieczkowski, G.A.G. Pereira, P.J.P.L. Teixeira, D.P.T. Thomazella

## Abstract

Witches’ broom disease (WBD) is a major constraint for cacao production in the Americas. The severe socioeconomic impact of WBD encouraged the evaluation of different control strategies, including the use of strobilurin fungicides. These molecules inhibit mitochondrial respiration, thus impairing ATP generation and leading to oxidative stress. These chemicals, however, have proven ineffective against the WBD pathogen *Moniliophthora perniciosa*. Here, we demonstrate that *M. perniciosa* tolerates high concentrations of strobilurins under *in vitro* conditions and highlight a set of molecular alterations that correlate with strobilurin tolerance in this fungus. Short-term exposure of *M. perniciosa* to the commercial strobilurin azoxystrobin led to the up-regulation of genes encoding enzymes of the glyoxylate cycle, gluconeogenesis, and fatty acid and amino acid catabolism, indicating that the fungal metabolism is remodeled to compensate for reduced ATP production. Furthermore, cell division, ribosome biogenesis, and sterol metabolism were repressed, which agrees with the impaired mycelial growth on azoxystrobin. Genes associated with cellular detoxification and response to oxidative stress (e.g., cytochrome P450s, membrane transporters and glutathione s-transferases) were strongly induced by the drug and represent potential strategies used by the pathogen to mitigate the toxic effects of the fungicide. Remarkably, exposure of *M. perniciosa* to azoxystrobin resulted in the spontaneous generation of a mutant with increased resistance to strobilurin. Comparative genomics and transcriptomics revealed alterations that may explain the resistance phenotype, including a large deletion in a putative transcriptional regulator and significant changes in the mutant transcriptome. Overall, this work provides important advances towards a comprehensive understanding of the molecular basis of strobilurin resistance in a tropical fungal pathogen. This is a fundamental step to efficiently employ these fungicides in agriculture and to prevent the emergence of strobilurin resistance.

## INTRODUCTION

The establishment and progress of human civilizations have been closely tied to the development of agriculture. Not surprisingly, plant diseases have been a main concern for humankind since ancient times (Santini et al. 2018). Major disease outbreaks, such as late blight, responsible for the Irish Potato Famine in the 19th century, or the Panama disease, which threatened the global banana production in the 1950s, have shown how plant pathogens can greatly impact our society, changing the course of human history (Yoshida et al. 2013; Ploetz et al. 2015). Currently, many other agronomically important crops are affected by pathogens, which cause significant yield losses and constraints to the global food supply (Rizzo et al. 2021). Thus, disease control is essential for ensuring efficient agricultural production.

We have battled the impact of pathogens in agriculture mostly through the cultivation of resistant plant varieties and/or the use of agrochemicals. However, continuous genetic changes within pathogen populations have often led to the loss of previously effective strategies for disease control. Resistant cultivars have historically been developed through selection and breeding programs. More recently, genetic manipulation based on biotechnological tools (i.e., genome editing technologies) has emerged as a promising alternative to classical breeding, which can be a time-consuming process. Yet, the use of genetic engineering to develop plants with improved agronomic traits is still subject to regulatory issues and public acceptance in many countries (Ahmad et al. 2021; Bearth et al. 2024). In addition to resistant varieties, agrochemicals have ensured an adequate food supply for many years and are particularly important for crops that are difficult to breed (e.g., banana, cassava). However, the indiscriminate use of these molecules can be highly harmful to both the environment and human health (Devi et al. 2022). Additionally, continued use of chemicals often selects for resistant/tolerant phytopathogens (Hu and Chen 2021).

The emergence of fungicide resistance in pathogen populations due to their extensive and long-term use is well-illustrated by strobilurin fungicides. Isolated from the basidiomycete *Strobilurus tenacellus*, strobilurin was modified for commercial use and promptly became one of the major classes of agricultural fungicides worldwide (Lamberth et al. 2021). The significant impact of these molecules on agriculture is exemplified by the synthetic strobilurin azoxystrobin, which is one of the most used fungicides in agriculture (Feng et al. 2020). Strobilurins belong to the quinol oxidation inhibitor (QoI) family, a class of molecules that inhibit mitochondrial respiration by specifically binding to the Qo domain of mitochondrial cytochrome bc_1_ (complex III) (Lamberth et al. 2021). These chemicals effectively reduced agricultural losses for several decades. However, one of their most prominent advantages, the highly precise mode of action, also proved to be their major limitation (Lamberth et al. 2021). Single-point mutations in the gene encoding cytochrome b (*Cytb*), which is part of the strobilurin-binding site at complex III, led to the emergence of fungal strains resistant to these highly specific inhibitors (Fernández-Ortuño et al., 2008).

Despite the effectiveness in controlling multiple plant pathogens, strobilurins have been largely ineffective against the witches’ broom disease (WBD) of cacao. WBD is one of the major phytopathological problems that impact the production of cocoa beans in the Americas and has caused devastating socioeconomic consequences, especially in Brazil (Teixeira et al. 2015). This disease is caused by the basidiomycete *Moniliophthora perniciosa,* and little is known about the molecular mechanisms involved in strobilurin tolerance in this fungus. The first insights on *M. perniciosa* tolerance to strobilurin were reported by Thomazella *et al*. (2012) with the identification of a single copy of an alternative oxidase gene (*Mp-Aox*) in the fungal genome. Alternative oxidase (AOX) is the main enzyme of the alternative respiratory pathway, and it limits the effectiveness of strobilurins *in planta* (Fernández-Ortuño, *et al*., 2008). It has been demonstrated that the Mp-AOX enzyme plays an important role in strobilurin tolerance (Thomazella et al. 2012). However, additional molecular mechanisms underpinning fungal tolerance to this class of fungicides have not been explored. Therefore, understanding the mechanisms associated with fungal response to these chemicals may lead to the identification of new molecular targets associated with strobilurin tolerance/resistance. Co-inhibition of these targets could completely halt fungal development and prevent the emergence of strobilurin-resistant strains, thus increasing fungicide efficiency and durability.

Here, we demonstrate that *M. perniciosa* tolerates unusually high concentrations of strobilurins *in vitro*, and this tolerance is not due to the typical mutations in the fungal *Cytb* gene. A time-course analysis of the fungal transcriptome following exposure to this fungicide indicated that the pathogen undergoes an intense metabolic reprogramming to mitigate the effects of the respiratory chain inhibition associated with reduced ATP production and increased oxidative stress. Many efflux pumps and detoxifying enzymes were up-regulated and may collectively contribute to fungal tolerance to strobilurin. Remarkably, long-term exposure of *M. perniciosa* mycelia to azoxystrobin led to the emergence of a resistant mutant with a highly distinct transcriptional signature when compared to wild-type isolates. Mutations leading to premature stop codons in two genes encoding putative growth and transcriptional regulators may have driven these massive alterations in the mutant transcriptome, contributing to an increased fungicide resistance. Taken together, our results provide valuable insights for the rational use and development of durable fungicide-based strategies to control the witches’ broom disease and possibly other important fungal diseases worldwide.

## METHODS

### Biological material

The *M. perniciosa* isolates FA553 and its derivatives FDS01 (FA553-derived sector 01; insensitive to azoxystrobin) and FDS02 (FA553-derived sector 02; parental phenotype) were used in this study. Fungal cultures were regularly maintained in Malt Yeast Extract Agar (MYEA) plates (17 g/L malt extract, 5 g/L yeast extract and 20 g/L agar) in an incubator at 28°C. Liquid cultures in Malt light medium (2 g/L malt extract, 5 g/L yeast extract and 50 mL/L glycerol) were maintained in Erlenmeyer flasks under 150 rpm agitation at 28°C.

### *M. perniciosa* mycelial growth in the presence of azoxystrobin

To evaluate the effects of long-term exposure of *M. perniciosa* to azoxystrobin, the fungicide Amistar WG (Syngenta; 50% of azoxystrobin and 50% inert compounds) was added to solid MYEA medium at a gradient of azoxystrobin concentrations: 1 mg/L, 10 mg/L, 20 mg/L, 50 mg/L, 100 mg/L, 200 mg/L and 500 mg/L. MYEA medium with no fungicide served as the control. Fungal growth was evaluated over a 30-day period, with photos and diameter measurements of each culture taken at regular intervals (7-, 14-, 21- and 28-days post inoculation).

### Evaluation of the *M. perniciosa* transcriptome in response to azoxystrobin

We evaluated the early effects of azoxystrobin on the *M. perniciosa* transcriptome through a time-course experiment spanning the first eight hours of exposure to the fungicide. Three biological replicates of each *M. perniciosa* isolate (FA553 and its derivatives FDS01 and FDS02) were cultivated separately in MYEA medium for 28 days. Fungal mycelia were then transferred to liquid cultures and incubated at 28°C under 150 rpm agitation. After seven days, two grams of wet mycelium were transferred to 20 mL of Malt light liquid medium, either without fungicide (control condition) or supplemented with 50 mg/L azoxystrobin (treated condition). Samples were harvested for RNA extraction at five time points: 0 minutes (i.e., immediately after fungal inoculation into the culture media), 30 minutes, 2 hours, 4 hours, and 8 hours. Supplementary Figure 1 summarizes the design of this time-course experiment.

### RNA extraction and RNA-seq library preparation and sequencing

RNA isolation was performed using the RNeasy Plant Kit (Qiagen) following the manufacturer’s instructions. RNA integrity and concentration were evaluated using the Lab Chip RNA high sensitivity assay (Caliper). A total of 90 RNA-seq libraries (30 for each isolate) were prepared from 1 µg of total RNA per sample, according to the TruSeq mRNA Stranded Sample Prep Kit protocol (Illumina). The Sciclone G3 Automated Liquid Handling Workstation (PerkinElmer) was used to prepare all libraries simultaneously. Quality control and final library concentrations were assessed using the DNA 1K Lab Chip high sensitivity assay (Caliper). Each individually barcoded library was combined into a single pool and sequenced across nine lanes of an Illumina HiSeq2500 instrument, generating an average of 13.9 million 50 bp single-end sequences per library. The RNA-seq data generated in this study is available at the NCBI Gene Expression Omnibus (GEO) under the accession number GSE281565.

### Read mapping and gene expression analysis

Quality of the raw reads was initially assessed using FASTX-Toolkit (Hannon 2010). Reads containing Illumina adapter sequences were identified and removed using Cutadapt. High-quality reads were then aligned to the *M. perniciosa* genome, obtained from the Witches’ Broom Genome Project (www.lge.ibi.unicamp.br/vassoura), using HiSAT2 (Kim et al. 2019). A maximum of one mismatch was allowed and reads mapping to multiple positions in the reference with equal alignment score were discarded. HTSeq (Anders et al. 2015) was used to count reads aligning to each of the 17,008 gene models. Differential expression analysis was performed with the edgeR package (Robinson et al. 2010), applying the False Discovery Rate (FDR) method (Benjamini and Hochberg 1995) for multiple-testing correction. To filter out weakly expressed genes, only those with a minimum expression level of 1 count per million in at least three libraries were included in the analysis. Normalization was performed using the trimmed mean of M-values method (TMM; function calcNormFactors in edgeR) (Robinson and Oshlack 2010). Count data were then fit into a negative binomial generalized linear model with a log link function. The experimental design followed a one-way layout and included ten groups representing the combinations of the two treatments in five time points. Contrasts were made between treated samples and controls at each time point and genes with FDR≤0.01 and a fold-change of at least 1.5x were considered differentially expressed.

### Hierarchical clustering and Principal Component Analyses

Principal Component Analysis (PCA) was conducted with the AMOR package in R (https://github.com/surh/AMOR), using the log_2_-transformed expression values (TPM, transcript per million) of the top 500 genes with the highest variance among samples. Hierarchical clustering analyses were performed with the ComplexHeatmap package in R (Gu et al. 2016). Gene expression values (TPM) were normalized by z-score transformation and clustered based on the Euclidean distance and the complete-linkage method.

### Functional classification of *M. perniciosa* genes

Functional annotations based on Gene Ontology (GO), InterPro (IPR) terms and TCDB classification (transporter classification database) were previously defined for each of the 17,008 *M. perniciosa* predicted genes (Teixeira et al. 2014). The annotation of transporters, glutathione transferases and cytochromes P450 were manually curated in this work. Enrichment analyses of GO and InterPro terms were conducted using the BinGO tool of Cytoscape (Maere et al. 2005). A significance threshold of FDR < 0.05 was used in these enrichment analyses.

### Identification of genomic variants in *M. perniciosa* genomes

To identify mutations potentially associated with the robust resistance to azoxystrobin in the FDS01 isolate, we employed a comprehensive pipeline for genomic variant calling in *M. perniciosa*. Whole-genome shotgun sequencing was performed on the FA553, FDS01, and FDS02 isolates using mycelial samples from the same cultures as those used in the RNA-seq experiment (Supplementary Figure 1). Additionally, an extra sample from the original FDS01 culture, along with its parental strain (FA553), were sequenced as temporally spaced replicates, providing further validation of the genomic data.

Genomic DNA was extracted following the protocol described by Mondego et al., (2008). DNA concentration was assessed using the Qubit DNA BR assay kit before library preparation. Libraries were constructed using 200 ng of genomic DNA with the TruSeq DNA Nano Kit (Illumina). Paired-end sequencing (2 × 100 bp) was conducted on the Illumina HiSeq 2500 platform at an average depth of 6.5 million paired-reads per sample, targeting approximately 30× coverage of the *M. perniciosa* genome.

Raw sequencing reads were subjected to quality control using FastQC v0.11.8 (Babraham Bioinformatics, UK). Adapter sequences and low-quality bases were trimmed using Trimmomatic v0.36 (Bolger et al. 2014), retaining only reads with a minimum length of 60 bp for further analysis. Clean reads were aligned to the *M. perniciosa* FA553 reference genome using Bowtie2 (Langmead and Salzberg 2012, 2) with default parameters. Genomic variants, including single nucleotide polymorphisms (SNPs) and small insertions/deletions (INDELs), were identified using two independent tools: FreeBayes v1.1.0 (Garrison and Marth 2012) and BCFtools mpileup v1.11 (Danecek et al. 2021). After marking duplicate reads using Picard Tools v2.23.4 (Broad Institute, USA), variant calling was performed for all five genomes. BCFtools isec v1.11 was used to identify variants unique to the FDS01 genome. Finally, variants called by FreeBayes were filtered using VcfFilter v0.2 with the following criteria: “QUAL > 20 & QUAL / AO > 10 & SAF > 1 & SAR > 1 & RPR > 1 & RPL > 1”. Variants identified by mpileup were filtered using SnpSift filter v4.3t with the criteria: “QUAL >= 20 && DP > 5 && MQ > 35”. The resulting variants were manually curated and categorized by their genomic location (exonic, intronic, intergenic) and their potential impact (e.g., synonymous, nonsynonymous, frameshift).

## RESULTS

### *M. perniciosa* tolerates high concentrations of strobilurins

Azoxystrobin is a synthetic strobilurin, with a recommended field dosage ranging from 40 mg/L to 120 mg/L of active ingredient (Amistar WG, Syngenta). We evaluated *M. perniciosa* growth in culture media containing increasing concentrations of this fungicide (ranging from 1 mg/L to 500 mg/L) over a 28-day period. While a clear reduction in the *M. perniciosa* growth rate was observed, none of the concentrations tested completely halted fungal development (Figure 1; Supplementary Figure 2). As expected, *M. perniciosa* growth was more severely affected at higher strobilurin concentrations. However, the fungus was still able to grow even at concentrations as high as 500 mg/L (Figure 1; Supplementary Figure 2).

**Figure 1.**
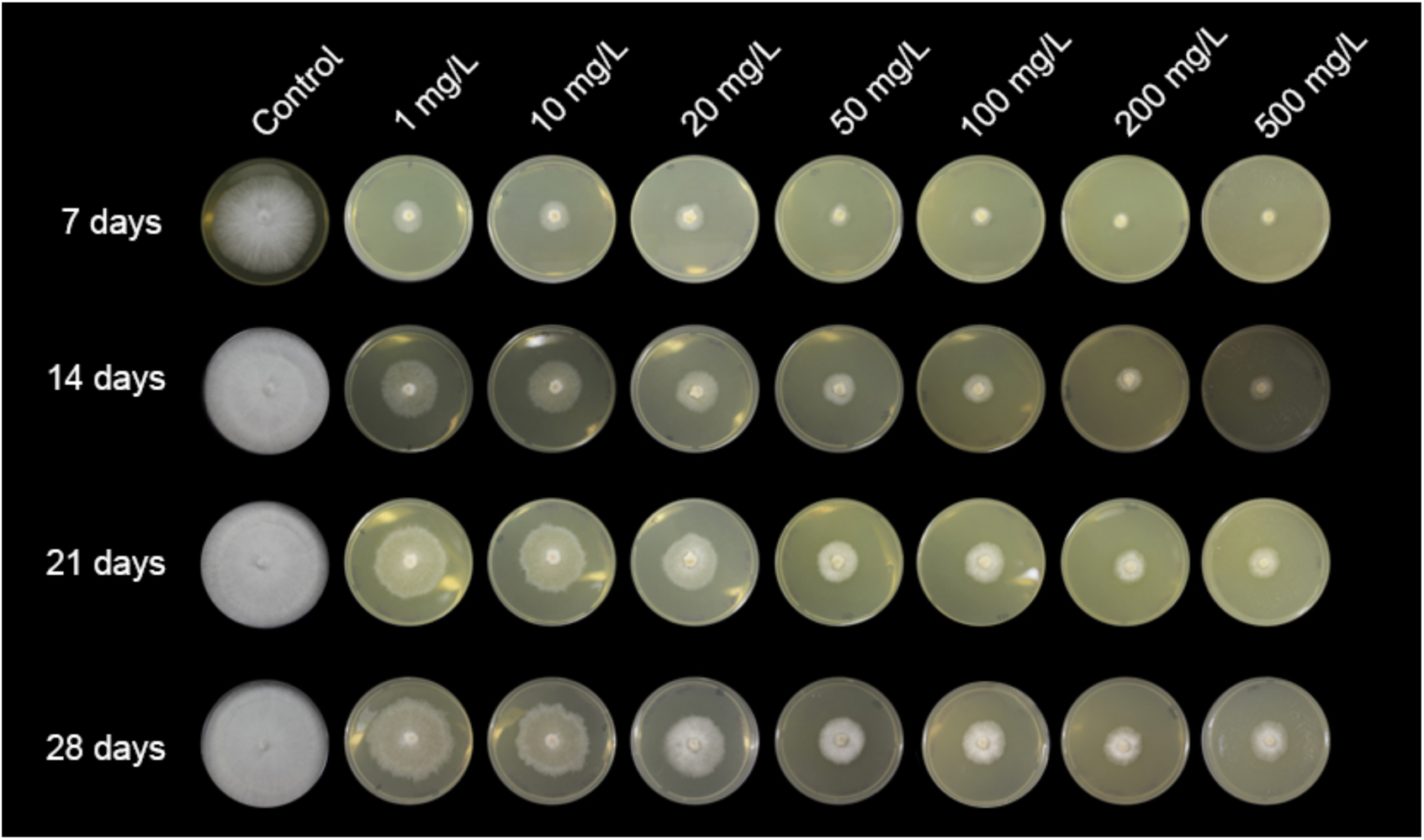
Evaluation of *M. perniciosa* growth in the presence of the strobilurin azoxystrobin. Cultivation of *M. perniciosa* under different concentrations of azoxystrobin for 28 days. Growth was observed at all tested concentrations. Although the fungal growth rate was affected in a dose-dependent manner, even concentrations as high as 500 mg/L were not sufficient to completely halt *M. perniciosa* growth. Representative images from three replicates are shown.

Interestingly, even after prolonged exposure (60 days) to the fungicide in solid media, *M. perniciosa* was able to recover its normal growth rate when transferred to media without fungicide, demonstrating that azoxystrobin treatment does not cause complete mycelial death. Furthermore, *M. perniciosa* also exhibited high tolerance to other two synthetic strobilurins (metominostrobin and picoxystrobin, Supplementary Figure 2). Remarkably, the *M. perniciosa* isolate used in this experiment (FA553) does not contain any of the typical mutations in the *CytB* gene that are known to confer resistance to strobilurins in other fungi (Supplementary Figure 3) (Sierotzki et al. 2000; Kim et al. 2003; Fernández-Ortuño et al. 2008). Therefore, other mechanisms may contribute to the high tolerance of this pathogen to this class of fungicides.

### Overview of the transcriptional responses of *M. perniciosa* to a strobilurin fungicide

To investigate the molecular responses associated with strobilurin tolerance in *M. perniciosa*, we performed a genome-wide gene expression analysis in a time-course experiment spanning the first eight hours following azoxystrobin exposure. The rationale was to capture the early response of the fungus to the stress caused by the drug. For the *M. perniciosa* isolate FA553, 30 RNA-seq libraries were constructed from mock and azoxystrobin-treated mycelium at five time points (0h, 0.5h, 2h, 4h and 8h). A total of 382.82 million Illumina reads were generated (Supplementary Figure 4A), with an average of 12.76 million reads per library (Supplementary Figure 4B), most of which mapped to exons (Supplementary Figure 4C). Principal Component Analysis (PCA) revealed a clear difference between the transcriptomes of control and azoxystrobin-treated mycelia (Figure 2A). Genes related to cellular detoxification and stress response contributed most to the separation between these two groups (described below).

**Figure 2.**
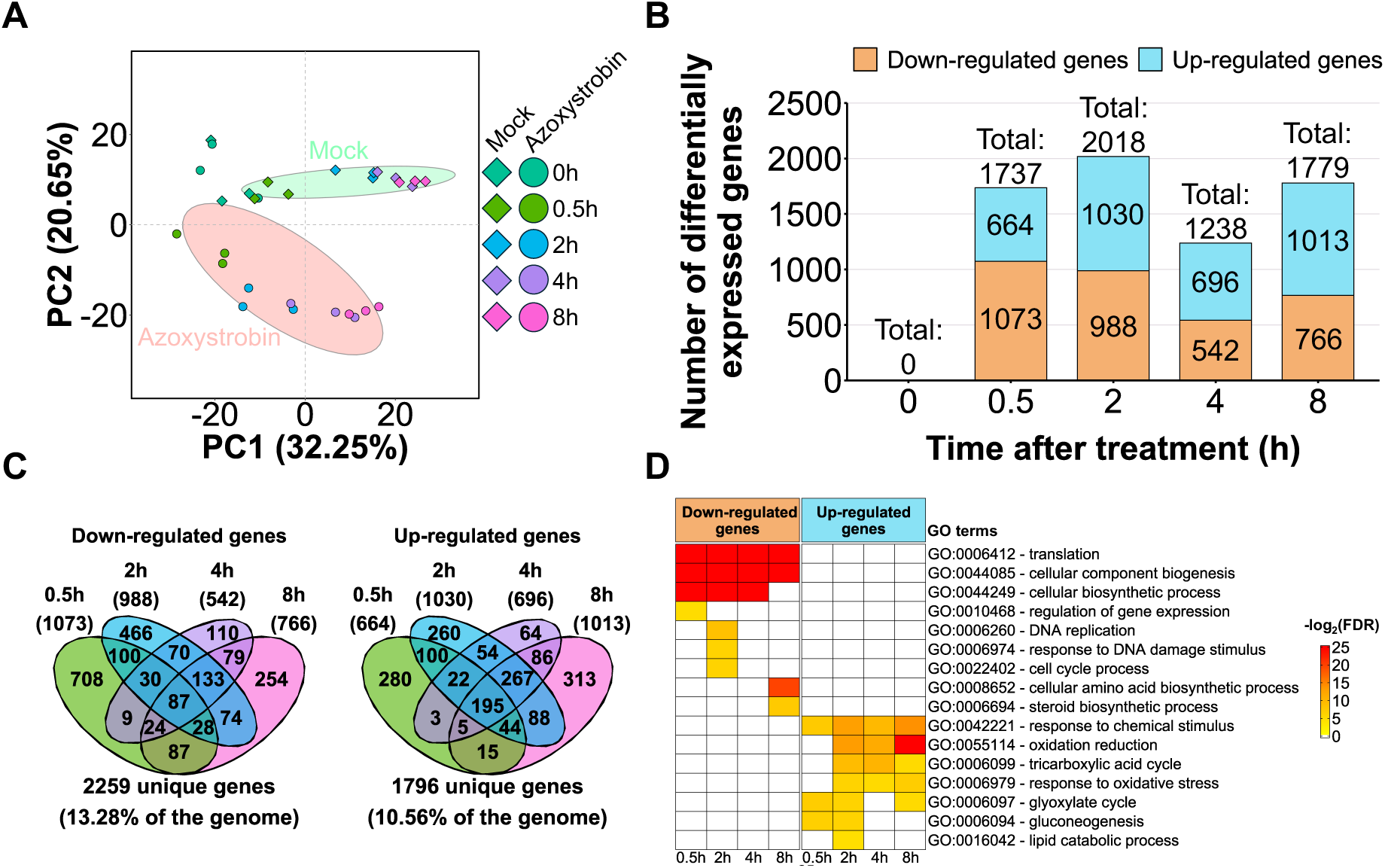
Overview of the transcriptional responses of *M. perniciosa* FA553 upon treatment with a strobilurin fungicide. (A) Principal Component Analysis (PCA) shows that the transcriptional profiles of mock and azoxystrobin-treated mycelia are clearly distinct at all time points, except for T=0h (samples harvested immediately after the start of the experiment). Ellipses show the parametric smallest area around the mean that contains 70% of the probability mass for each group. (B) Number of differentially expressed genes at each time point of the experiment (0h, 0.5h, 2h, 4h and 8h). Here, azoxystrobin-treated mycelia were compared to their corresponding mock-treated controls. (C) Venn diagrams demonstrating the overlap of DEGs across different time points (left panel: down regulated genes, right panel: up regulated genes). (D) Gene ontology (GO) enrichment analysis revealed the up-regulation of catabolic processes and energy metabolism pathways and down-regulation of biological processes related to cellular multiplication, DNA replication and protein synthesis at most time points. The complete GO enrichment analysis and the corresponding statistics are provided in Supplementary Data Set 2.

Differentially expressed genes between mock and azoxystrobin-treated samples (false discovery rate [FDR] ≤ 0.01; fold-change of 1.5) were identified using edgeR (Robinson et al. 2010). Of the 17,008 gene models in the *M. perniciosa* isolate FA553, 14,399 were considered expressed in our experiment (see Methods for details). Among these, 3,969 were differentially expressed in at least one time-point of the experiment, representing 23% of the total *M. perniciosa* genes (Supplementary Data Set 1). As expected, no genes were differentially expressed in samples harvested immediately after transferring the mycelium to azoxystrobin-amended culture medium (T=0h). The number of differentially expressed genes (DEGs) at subsequent time points varied from 1,238 (T=4h) to 2,018 (T=2h) (Figure 2B). Importantly, as little as 30 min of exposure to azoxystrobin was sufficient to affect the expression of thousands of genes. Overall, fungal responses to azoxystrobin at different time points were highly similar, with 1,660 genes (41.82% of DEGs) differentially expressed at more than one time point (Figure 2C; Supplementary Data Set 1). Interestingly, gene ontology (GO) enrichment analysis revealed intense metabolic reprogramming, including up-regulation of biological processes related to energy metabolism, such as cellular respiration, gluconeogenesis, and lipid catabolism (Figure 2D; Supplementary Data Set 2). Conversely, biological processes associated with growth, such as DNA replication, translation, and cell cycle, were down-regulated.

### The *M. perniciosa* respiratory chain is remodeled in response to azoxystrobin

In agreement with our previous studies (Thomazella et al. 2012; Barsottini et al. 2019; Moretti-Almeida et al. 2019), *M. perniciosa* seems to mitigate the effects of mitochondrial complex III inhibition by the rapid up-regulation of the *Mp*-*Aox* gene (Figure 3A), resulting in the activation of an alternative respiratory pathway that is insensitive to strobilurin. Indeed, AOX enzymatic activity has only been detected in *M. perniciosa* upon azoxystrobin exposure (Thomazella et al. 2012). Because this alternative pathway is less energy-efficient than the main respiratory chain (Luévano-Martínez et al. 2019), *M. perniciosa* cells likely experience ATP deprivation when exposed to azoxystrobin. Under these conditions, other enzymes and pathways may also play important roles to ensure fungal survival. A gene encoding a non-proton pumping internal NADH dehydrogenase (NDH-2) was differentially expressed 2h after treatment (Figure 3A), possibly to alleviate azoxystrobin-induced oxidative stress. Additionally, at least 12 genes encoding components of the main mitochondrial respiratory chain (including subunits of complexes II, III and IV, and ATP synthase) were induced by azoxystrobin (Figure 3B; Supplementary Data Set 3), likely as an attempt to compensate for the impaired cellular respiration. These results suggest that azoxystrobin treatment might cause the rearrangement and assembly of additional respiratory complexes (Supplementary Data Set 3) to compensate for the decreased energy production caused by complex III inhibition.

**Figure 3.**
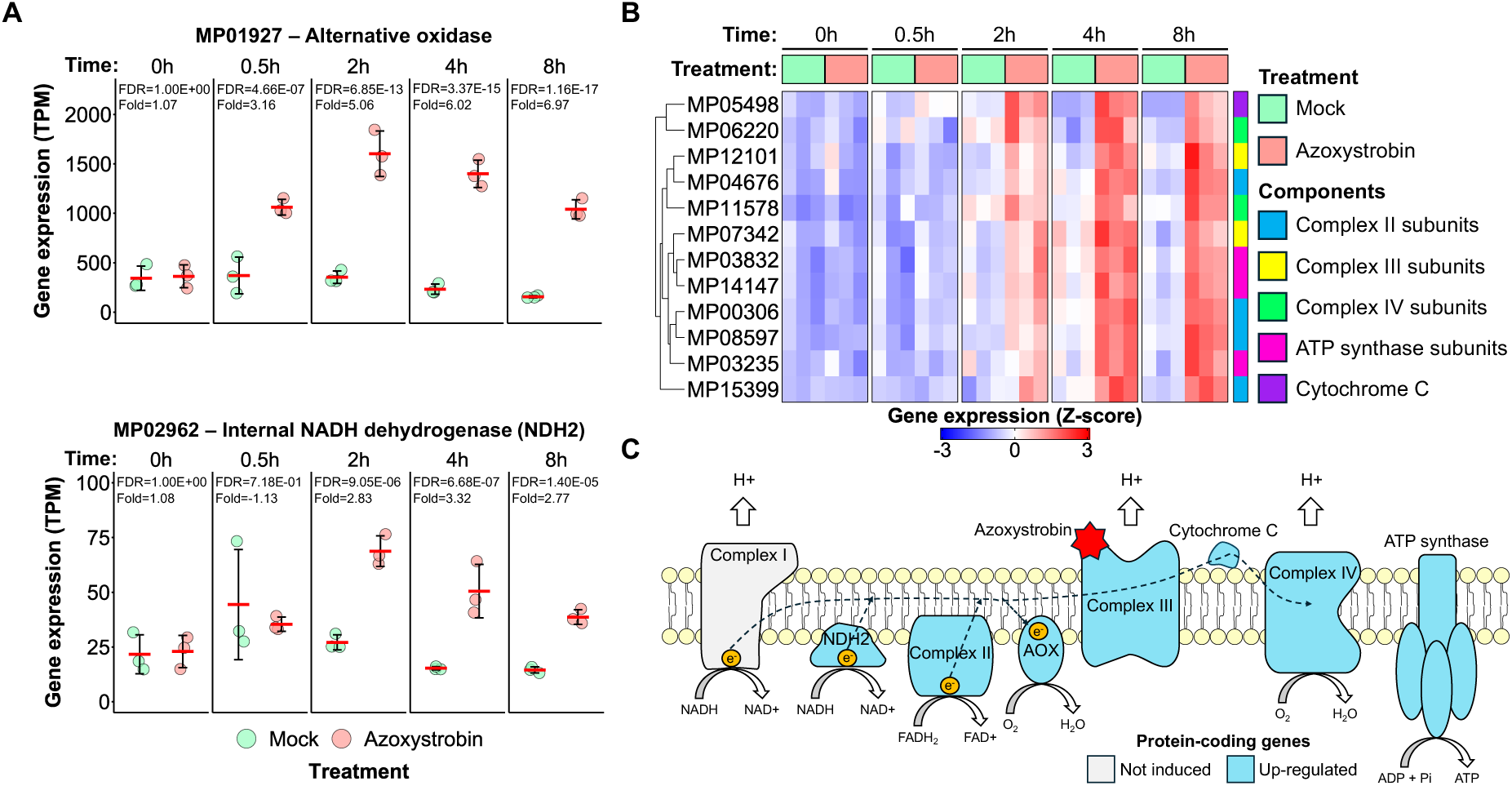
Azoxystrobin induces genes encoding components of the mitochondrial respiratory chain. (A) Genes coding for an alternative oxidase (MP01927) and an internal NADH dehydrogenase (MP02962) are significantly up-regulated upon azoxystrobin exposure. (B) Expression pattern of genes coding for components of the main mitochondrial respiratory chain. Note the induction of components of most mitochondrial respiratory complexes, possibly as a compensatory response to alleviate the negative effects of azoxystrobin on ATP production. The complete list of differentially expressed genes associated with cellular respiration (including their functional annotation) is provided in Supplementary Data Set 3. (C) Schematic representation of the mitochondrial respiratory chain, highlighting proteins whose corresponding genes were upregulated upon azoxystrobin treatment.

### Azoxystrobin induces fatty acid and amino acid catabolism and the TCA cycle

In the presence of azoxystrobin, *M. perniciosa* seems to activate the catabolism of lipids to boost the supply of reduced coenzymes for the respiratory chain (Supplementary Data Set 3). GO enrichment analysis revealed that ‘lipid catabolic processes’ (GO:0016042) were activated after 2h of fungal exposure to azoxystrobin (Figure 2D). Indeed, a total of 18 genes involved in fatty acid catabolism were differentially expressed in the experiment (Supplementary Data Set 3). Six genes encoding lipases, which catalyze the hydrolysis of triacylglycerols into free fatty acids and glycerol, were up-regulated by azoxystrobin (Supplementary Data Set 3). Furthermore, genes required for fatty acid activation (acyl-CoA synthase and acyl-carnitine transferase) were also strongly activated in azoxystrobin-treated mycelia (Supplementary Data Set 3). Consistent with the use of fatty acids as an energy source, genes encoding three mitochondrial enzymes involved in β-oxidation (enoyl-CoA hydratase, 3-hydroxyacyl-CoA dehydrogenase, and 3-ketoacyl-CoA thiolase) were up-regulated in response to the fungicide (Supplementary Data Set 3). Notably, β-oxidation of fatty acids also occurs in peroxisomes (Poirier et al. 2006). Accordingly, genes encoding the peroxisomal enzymes acyl-CoA oxidase, hydratase-dehydrogenase-epimerase and 3-ketoacyl thiolase, were also induced (Supplementary Data Set 3). Thus, both peroxisomal and mitochondrial β-oxidation pathways were activated in response to azoxystrobin.

Reduced coenzymes (NADH and FADH_2_) and acetyl-CoA are the final products of β- oxidation. Both reduced coenzymes can serve as electron donors in the mitochondrial electron transport chain for ATP production (Letts and Sazanov 2017). Moreover, acetyl-CoA can enter the tricarboxylic acid cycle (TCA) (Figure 4), generating even more NADH, FADH_2_ and ATP (Akram 2014). The degradation of branched-chain amino acids (valine, leucine and isoleucine) also contributes to ATP production by providing the TCA cycle with acetyl-CoA (Hildebrandt et al. 2015). Notably, 9 genes encoding enzymes required for the catabolism of branched-chain amino acids were induced by azoxystrobin (Supplementary Data Set 3). Interestingly, the GO term ‘Tricarboxylic Acid Cycle’ (GO:0006099) was enriched in the set of genes up-regulated by the fungicide (Figure 2D). In particular, genes encoding three of the enzymes that catalyze the steps of the TCA cycle where the coenzymes NAD^+^ and FAD are reduced (i.e., α-ketoglutarate dehydrogenase, malate dehydrogenase and succinate dehydrogenase) were all activated in azoxystrobin-treated mycelia (Figure 4). These findings are consistent with the hypothesis that lipids and amino acids are catabolized to sustain cellular respiratory capacity and energy production upon azoxystrobin treatment, thus contributing to fungal survival.

**Figure 4.**
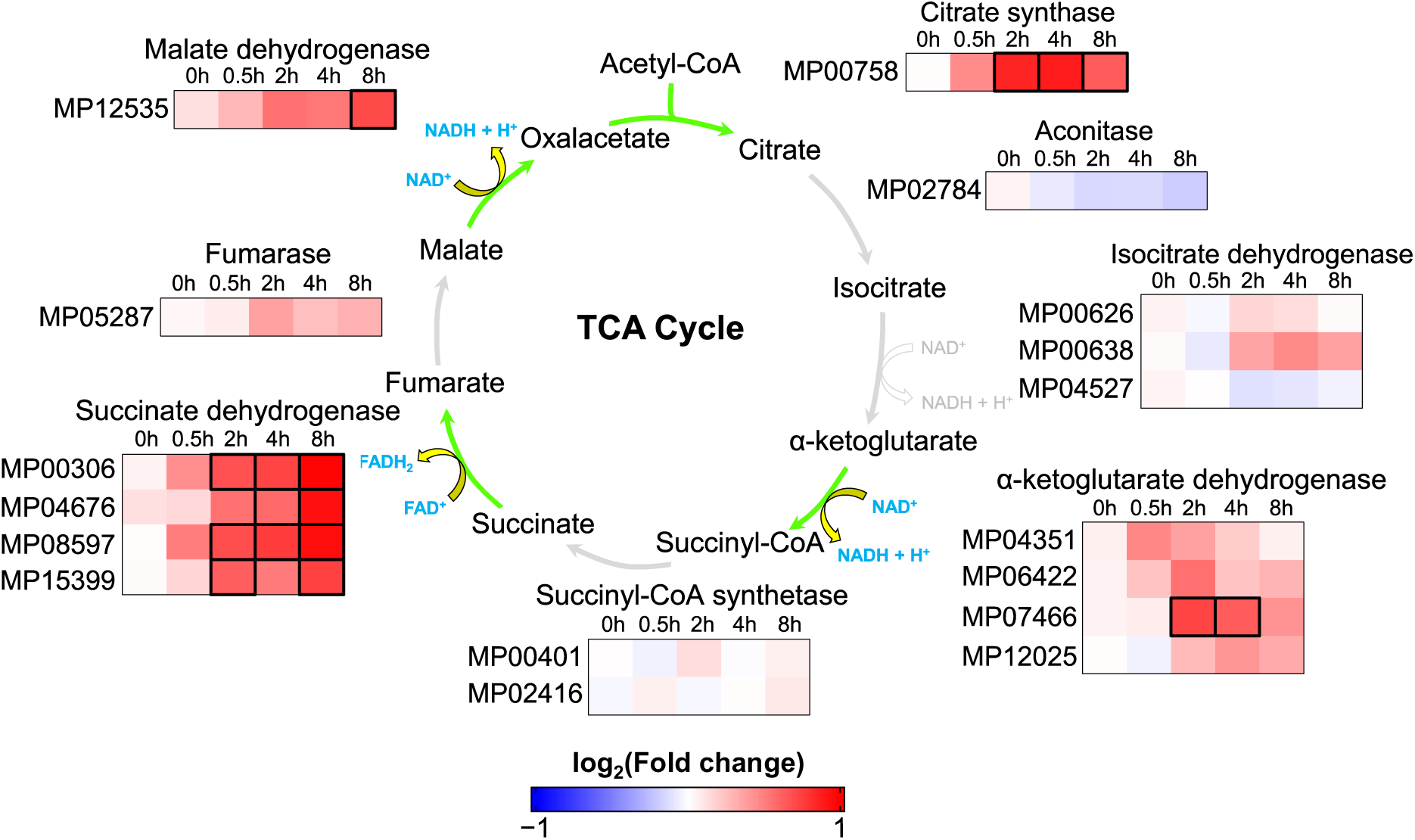
Key components of the TCA cycle are induced in azoxystrobin-treated mycelia. The expression pattern of genes encoding enzymes of the TCA cycle are shown. Green arrows indicate azoxystrobin-induced reactions, while the reduced coenzymes produced in these reactions are highlighted in blue. Differentially expressed genes at specific time points are indicated by black bold borders. The complete gene expression data and differential expression results are provided in Supplementary Data Set 3.

### The glyoxylate cycle and gluconeogenesis are activated by azoxystrobin

In fungi, fatty acids can also be converted into glucose and precursors of the TCA cycle via the glyoxylate cycle (Chen et al. 2016). This process occurs in glyoxysomes, which are specialized peroxisomes that contain the enzymes isocitrate lyase and malate synthase, both exclusive of the glyoxylate cycle (Pracharoenwattana and Smith 2008). Remarkably, genes encoding these enzymes are induced at three (isocitrate lyase) and four (malate synthase) different time points (Supplementary Data Set 3). Isocitrate lyase cleaves isocitrate into succinate and glyoxylate. Glyoxylate is then condensed with acetyl-CoA by malate synthase, producing malate, whereas succinate is transported to the mitochondrial matrix through a dicarboxylate transporter (Palmieri and Monné 2016). In agreement, a mitochondrial dicarboxylate transporter is activated after 2 hours of azoxystrobin treatment (Supplementary Data Set 3). Succinate then enters the TCA cycle, where it is converted into malate, which is transported to the cytosol and converted into glucose via enzymes involved in gluconeogenesis, such as PEPCK (Phosphoenolpyruvate carboxykinase), fructose 1,6- bisphosphatase and glucose 6-phosphatase. Except for glucose 6-phosphatase, which catalyzes the conversion of glucose 6-phosphate to glucose, genes encoding these enzymes were activated at all four time points (Supplementary Data Set 3). These results indicate that both the glyoxylate cycle and gluconeogenesis pathways are induced in response to azoxystrobin.

### Fitness costs associated with azoxystrobin exposure

*M. perniciosa* shows reduced growth in the presence of azoxystrobin (Figure 1; Supplementary Figure 2). In agreement, biological processes associated with cell growth and proliferation were repressed in response to the fungicide (Figure 2D). Genes with GO annotation for cell cycle (GO:0022402), DNA replication (GO:0006260), gene expression (GO:0010468), translation (GO:0006412) and steroid biosynthesis (GO:0006694) were repressed upon azoxystrobin treatment (Figure 2D). A total of 358 genes related to these processes were down-regulated in, at least, one time point of the experiment (Supplementary Data Set 4). These results suggest that basic biological processes are suppressed in response to energy-depriving conditions triggered by azoxystrobin.

#### Cell cycle progression and DNA replication

A set of 29 genes required for cell cycle progression was repressed by azoxystrobin treatment (Supplementary Data Set 4), including those encoding the cdk2-cdc13 complex and the M-phase inducer phosphatase, which are essential for cells to enter mitosis (Kimelman 2014; Gutiérrez-Escribano and Nurse 2015). In addition, 46 genes coding for DNA primases, polymerases, ligases, and components of the pre-replication complex, which are important to initiate DNA replication (O’Donnell et al. 2013), were also repressed, mostly, at the 2h-time point (Supplementary Data Set 4). Remarkably, we verified down-regulation of 13 genes associated with DNA repair, such as the DNA polymerase β, RAD52 and PCNA (Lisby and Rothstein 2009; Yamtich and Sweasy 2010; Boehm et al. 2016) (Supplementary Data Set 4). Collectively, these results suggest that both the cell cycle and DNA replication/repair processes are impaired in response to azoxystrobin.

#### mRNA and protein synthesis

A total of 61 genes coding for components of the transcriptional machinery were repressed in response to azoxystrobin in at least one time point (Supplementary Data Set 4). These include the gene coding for Rpa49, a subunit of RNA polymerase I (Albert et al. 2011) and CDC73 (Cell Division Control 73), which is a constituent of the PAF1 complex, a key regulator of mRNA synthesis (Van Oss et al. 2017). Furthermore,193 genes coding for translation-related proteins were mostly repressed 30 minutes after azoxystrobin exposure (Supplementary Data Set 4). They include 80 genes encoding components of the 40S and 60S ribosomal subunits and 18 genes coding for tRNA synthetases. These results indicate that both the transcription and translation machineries of *M. perniciosa* are negatively affected by azoxystrobin treatment.

#### Ergosterol biosynthesis

Of the 23 genes putatively involved in ergosterol biosynthesis (Jordá and Puig 2020), 16 were down-regulated in mycelia exposed to azoxystrobin in at least one time point (Supplementary Data Set 4). These include genes encoding the enzymes hydroxymethylglutaryl-CoA synthase, lanosterol 14-α-demethylase and δ-sterol reductase (Supplementary Data Set 4), all of which are necessary for ergosterol biosynthesis. Notably, ergosterol is a major component of fungal cell membranes, playing a role in maintaining membrane fluidity and integrity (Jordá and Puig 2020). Our results suggest that azoxystrobin interferes with ergosterol biosynthesis, potentially compromising membrane integrity and impairing fungal growth.

#### *M. perniciosa* activates a variety of detoxification mechanisms in response to azoxystrobin

In addition to the metabolic rearrangement observed in *M. perniciosa* in response to azoxystrobin, the analysis of the fungal transcriptome suggested the activation of diverse detoxification mechanisms in the azoxystrobin-treated mycelium (Figure 5A; Supplementary data set 5). These mechanisms may independently function to modify, degrade, or excrete the fungicide from fungal cells, alleviating its toxic effects. Moreover, genes associated with the mitigation of oxidative stress were also up-regulated in the azoxystrobin-treated mycelium.

**Figure 5.**
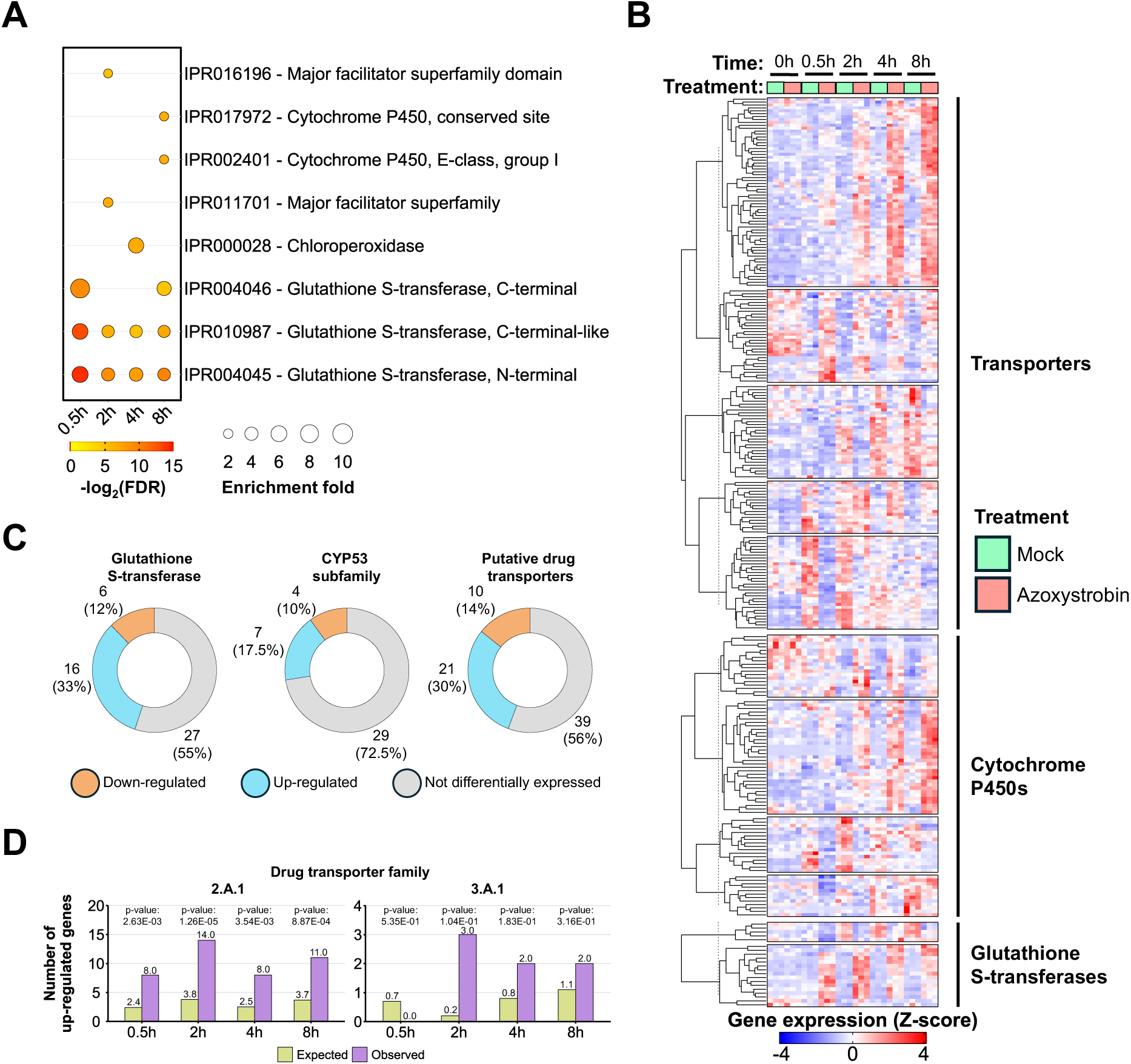
Activation of detoxification mechanisms in *M. perniciosa* in response to azoxystrobin. (A) InterPro enrichment analysis showing overrepresented terms in the azoxystrobin-treated mycelium (FDR < 0.05). (B) Hierarchical clustering of genes encoding detoxification factors that are differentially expressed upon treatment with azoxystrobin. (C) Number of detoxification factors up-regulated, down-regulated or not differentially expressed in the presence of azoxystrobin. (D) Enrichment analysis of candidate drug transporters in the 2.A.1 and 3.A.1 superfamilies (MFS and ABC transporters, respectively). Members of the 2.A.1 superfamily are more prevalent among the set of up-regulated genes than the expected by chance. Genes encoding members of the 3.A.1 superfamily are activated by azoxystrobin, but the enrichment of this superfamily is not statistically significant.

#### Glutathione S-transferase (GST)

GSTs play a central role in drug resistance and detoxification pathways. These enzymes catalyze the conjugation of glutathione to toxic endogenous or xenobiotic substrates, preventing these molecules from reacting with essential cellular macromolecules (Gullner et al. 2018). The *M. perniciosa* genome encodes 52 GST genes, of which 15 (29%) were induced and only 6 (11%) were repressed by azoxystrobin, suggesting a role for glutathione metabolism in the *M. perniciosa* tolerance to strobilurin fungicides (Figure 5B and 5C; Supplementary data set 5).

#### Cytochrome P450

Members of the cytochrome P450 (CYP450) family were also differentially expressed in the presence of azoxystrobin. CYP450 enzymes can catalyze the conversion of xenobiotic compounds into easily excreted, less toxic, products. Interestingly, the CYP450 family is expanded in the *M. perniciosa* genome (Mondego et al. 2008), suggesting a high potential for detoxification in this fungus. Of the 326 CYP450 genes in *M. perniciosa*, 79 (24%) were differentially expressed (52 up-regulated, 27 down-regulated) (Figure 5B; Supplementary data set 5). In addition, 23 genes coding for CYP450s in *M. perniciosa* belong to the CYP53 subfamily, which is involved in the detoxification of benzoate, an intermediate in the degradation of aromatic compounds in fungi (Harwood and Parales 1996). Interestingly, five of these genes were induced by azoxystrobin (Figure 5C; Supplementary data set 5), which is an aromatic compound, suggesting the participation of CYP53 in azoxystrobin degradation.

#### Efflux pumps

At least 179 genes encoding members of different families of transporters were differentially expressed in response to azoxystrobin. The *M. perniciosa* genome encodes 1,413 putative transporters, including 258 from the 2.A.1 superfamily (<u>M</u>ajor <u>F</u>acilitator <u>S</u>uperfamily - MFS) and 56 from the 3.A.1 superfamily (<u>A</u>TP-<u>b</u>inding <u>c</u>assette - ABC) (Figure 5D; Supplementary data set 5). These transporters likely function as efflux pumps, transporting xenobiotic compounds by transporting them to the extracellular space and preventing their lethal accumulation (Ghannoum and Rice 1999). Within these subfamilies, at least 70 transporters were classified as putative drug transporters, and 21 of them (17 of the 2.A.1 and 4 of the 3.A.1 superfamily) were up-regulated by azoxystrobin (Figures 5D and 5E; Supplementary data set 5). These findings suggest that *M. perniciosa* uses membrane transporters to pump azoxystrobin to the extracellular space, thus preventing its toxic accumulation in the fungal cell.

### Exposure of *M. perniciosa* to azoxystrobin promoted the emergence of a resistant mutant

The transcriptional analysis of *M. perniciosa* in response to azoxystrobin revealed several molecular alterations that may enable fungal survival in the presence of this fungicide, albeit at the cost of reduced growth. Remarkably, during the cultivation of *M. perniciosa* isolate FA553 on solid medium containing azoxystrobin (200 mg/L), we noticed the development of a morphologically distinct mycelial sector that grew faster than the rest of the mycelium on the same plate (Figure 6A). This presumably resistant mycelial sector was isolated, replicated *in vitro* and named FDS01 (FA553-derived sector 01). Another sector from the same Petri dish, displaying wild-type-like morphology, was also isolated and named FDS02 (FA553-derived sector 02). The three isolates (i.e., FA553, FDS01 and FDS02) were subsequently cultivated in the presence of azoxystrobin to evaluate their sensitivity to this fungicide (Figure 6B). As expected, the growth of the FA553 and FDS02 isolates was inhibited, but not entirely blocked by azoxystrobin. Remarkably, FDS01 showed vigorous growth even at high concentrations of azoxystrobin (i.e., 500 mg/L), suggesting that this isolate is resistant to the fungicide (Figure 6B).

**Figure 6.**
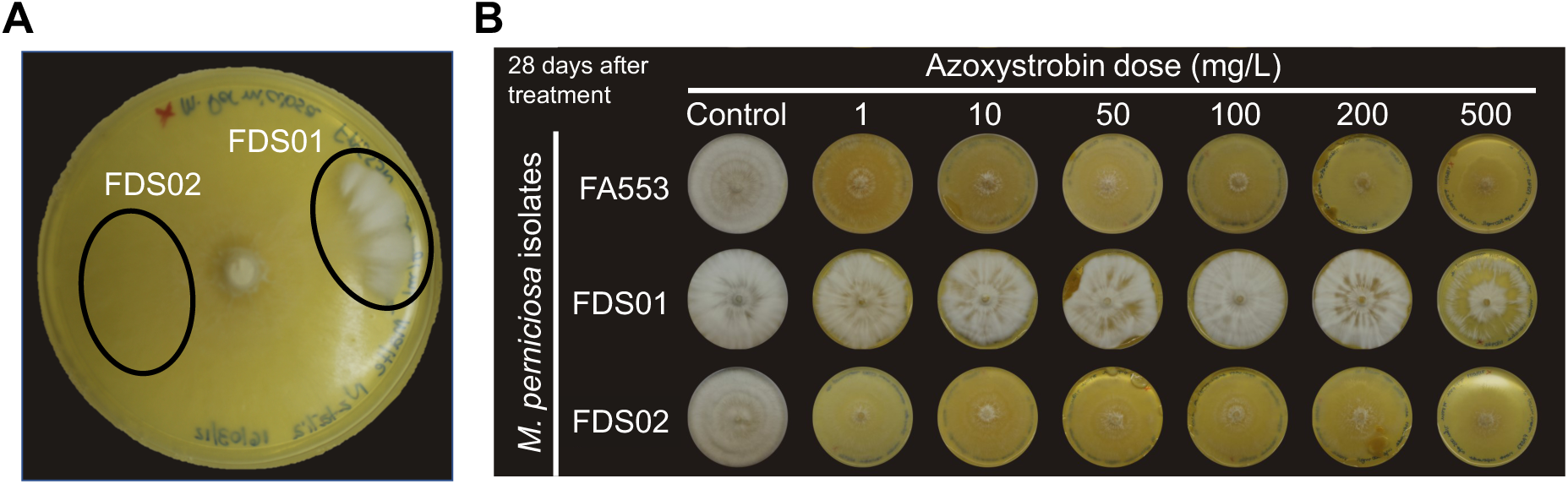
Isolation of an *M. perniciosa* mutant with reduced sensitivity to azoxystrobin. (A) In the presence of azoxystrobin, *M. perniciosa* grows as thin mycelia on solid medium. Interestingly, a mycelial sector with an increased growth rate was observed, indicating the emergence of a phenotype with enhanced resistance to the fungicide. This mycelial sector was isolated and named FDS01. As a control, a sector displaying wild-type morphology was also isolated and named FDS02. The image was taken 41 days after inoculation. (B) The isolates FDS01, FDS02 and FA553 (the parental isolate) were cultivated in media with increasing concentrations of azoxystrobin. All three isolates display similar morphology in MYEA medium. However, while the isolates FA553 and FDS02 grow as thin mycelia in the presence of azoxystrobin, the FDS01 isolate shows vigorous growth, even at higher fungicide concentrations. Images were taken 28 days after inoculation.

As described above, single-point mutations in the *CytB* gene can confer resistance to strobilurin fungicides (Fernández-Ortuño et al. 2008). These mutations, such as F129L and G143A, alter the binding of strobilurins to cytochrome B, abolishing their fungicidal effect. However, no such mutation in the cytochrome B gene was detected in the FDS01 mutant (Supplementary Figure 3). Additionally, although AOX has already been shown to mitigate azoxystrobin effects on *M. perniciosa* (Thomazella et al. 2012), *Mp*-*Aox* is similarly up-regulated in all three isolates along the time-course experiment (Supplementary Figure 5). This suggests that the azoxystrobin-resistant phenotype of the FDS01 isolate is unlikely to be due to differences in Mp-AOX activity levels. These results indicate that well-described strobilurin resistance mechanisms (i.e., mutations in the *CytB* gene and Mp-AOX activity) do not account for the increased tolerance of the FDS01 isolate to azoxystrobin, implying the involvement of additional mechanisms.

### The FDS01 isolate shows intense transcription deregulation

The RNA-seq results described above focus on the FA553 isolate, highlighting the global effects of azoxystrobin on the wild-type mycelia of *M. perniciosa*. To investigate possible transcriptional changes linked to the azoxystrobin resistance phenotype of FDS01, RNA-seq experiments were also conducted on the FDS01 and FDS02 isolates. Along with the 30 RNA-seq libraries for FA553, 60 additional libraries were generated for FDS01 and FDS02 (30 for each isolate; Supplementary Figure 4). These libraries correspond to biological triplicates of mock and azoxystrobin-treated samples at five time points (i.e., 0h, 0.5h, 2h, 4h and 8h).

A Principal Component Analysis (PCA) revealed that the parental strain FA553 and its wild-type derivative FDS02 share a highly similar transcriptional signature (Figure 7A). This result validates the key conclusions regarding the effects of azoxystrobin on *M. perniciosa* using a second isolate (FDS02).Conversely, the azoxystrobin-resistant mutant FDS01 was clearly different from the other two isolates, regardless of the presence of azoxystrobin. Yet, the fungicide had a pronounced effect on the transcriptomes of all three genotypes throughout the time course (Figure 7A), supporting the observation that azoxystrobin triggers extensive transcriptional reprogramming of fungal metabolism.

**Figure 7.**
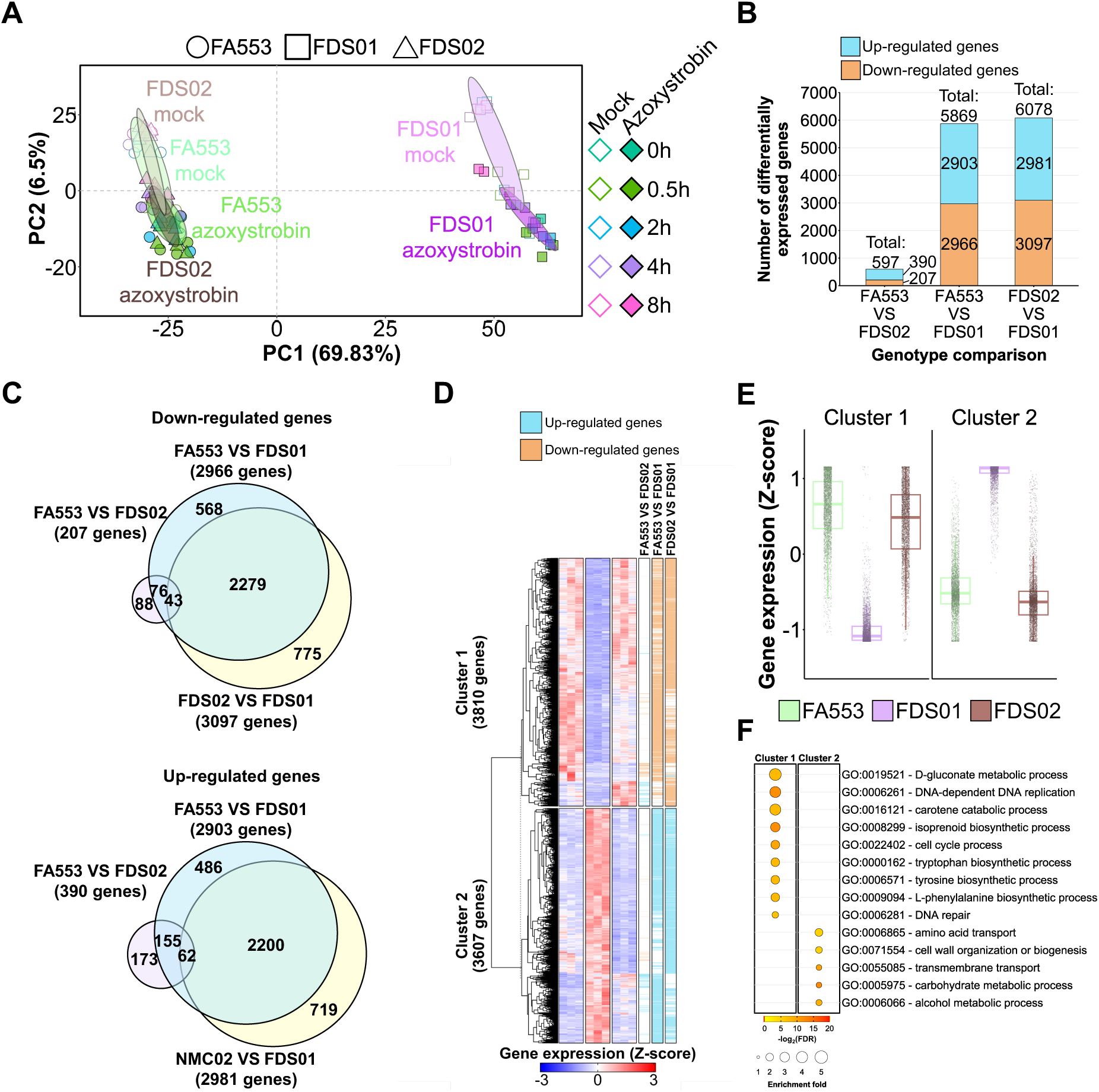
The mutant strain FDS01 shows an altered transcriptional program. (A) Principal Component Analysis (PCA) plot based on the transcriptomes of the isolates FA553, FDS01, and FDS02. The FDS01 transcriptional profile is remarkably different from the transcriptomes of the other two isolates, regardless of the presence of azoxystrobin. The color code represents different time points in the experiment. Empty diamonds represent control samples, while filled diamonds represent azoxystrobin-treated samples of the three isolates. Circles, squares, and triangles represent triplicates from FA553, FDS01 and FDS02, respectively. Ellipses show the parametric smallest area around the mean that contains 60% of the probability mass for each group. (B) Number of down-regulated (orange) and up-regulated (blue) genes between genotypes. (C) Overlap of down-regulated (top) and up-regulated genes (bottom) based on the comparison of the genotypes at time point T=0h. (D) Hierarchical clustering of the 7417 differentially expressed genes among the genotypes FA553, FDS01 and FDS02. The FDS01 mutant displays a distinct transcriptional signature compared to the FA553 and FDS02 wild-type isolates. (E) Expression patterns for clusters 1 and 2, as defined in the heatmap shown in panel D. (F) Gene Ontology (GO) enrichment analysis indicates the enriched biological processes in each cluster. A detailed GO enrichment analysis, including the corresponding statistics, can be found in Supplementary Data Set 7.

To further explore the differences between FDS01 and the other two isolates, we performed pairwise comparisons in the absence of azoxystrobin at time point T=0h. A total of 7,417 genes were differentially expressed (FDR ≤ 0.01 and Fold-change ≥ 1.5) across the three genotypes (Supplementary Data Set 6). The FDS01 mutant showed 5,869 and 6,078 differentially expressed genes compared to FA553 and FDS02, respectively (Figure 7B, C), whereas only 597 genes were differentially expressed between FA553 and FDS02. These results confirm the high similarity between FA553 and FDS02, while underscoring the distinct nature of the FDS01 mutant.

Hierarchical clustering of the differentially expressed genes among genotypes revealed two distinct clusters (Figure 7D). Cluster 1 consists of genes that were more highly expressed in the wild-type genotypes (FA553 and FDS02), but weakly expressed in the FDS01 mutant (Figure 7D and 7E). Conversely, cluster 2 contains genes with higher expression in FDS01 and lower expression levels in the other two isolates. Notably, cluster 1 includes genes involved in basic biological processes, such as DNA replication, cell cycle and amino acid biosynthesis, while cluster 2 is enriched for genes associated with metabolic process, transmembrane transport, and cell wall biogenesis (Figure 7F; Supplementary Data Set 7).

### Putative growth and transcriptional regulators are mutated in FDS01

A genomic survey was conducted to identify differences in FDS01 compared to the wild-type genotypes FA553 and FDS02. For this, genomic DNA from these isolates was shotgun-sequenced, and the resulting reads were aligned to the *M. perniciosa* reference assembly (isolate FA553). Four genomic differences were consistently identified by the two variant-calling methods employed (freebayes and Mpileup) in FDS01, but not in FA553 and FDS02 (Figure 8A and 8B; Supplementary Data Set 8). Three of these variants were single nucleotide polymorphisms (SNPs) in protein-coding genes (MP02676, MP10135 and MP12249), whereas the remaining one was an 8-nucleotide deletion in MP01179.

**Figure 8.**
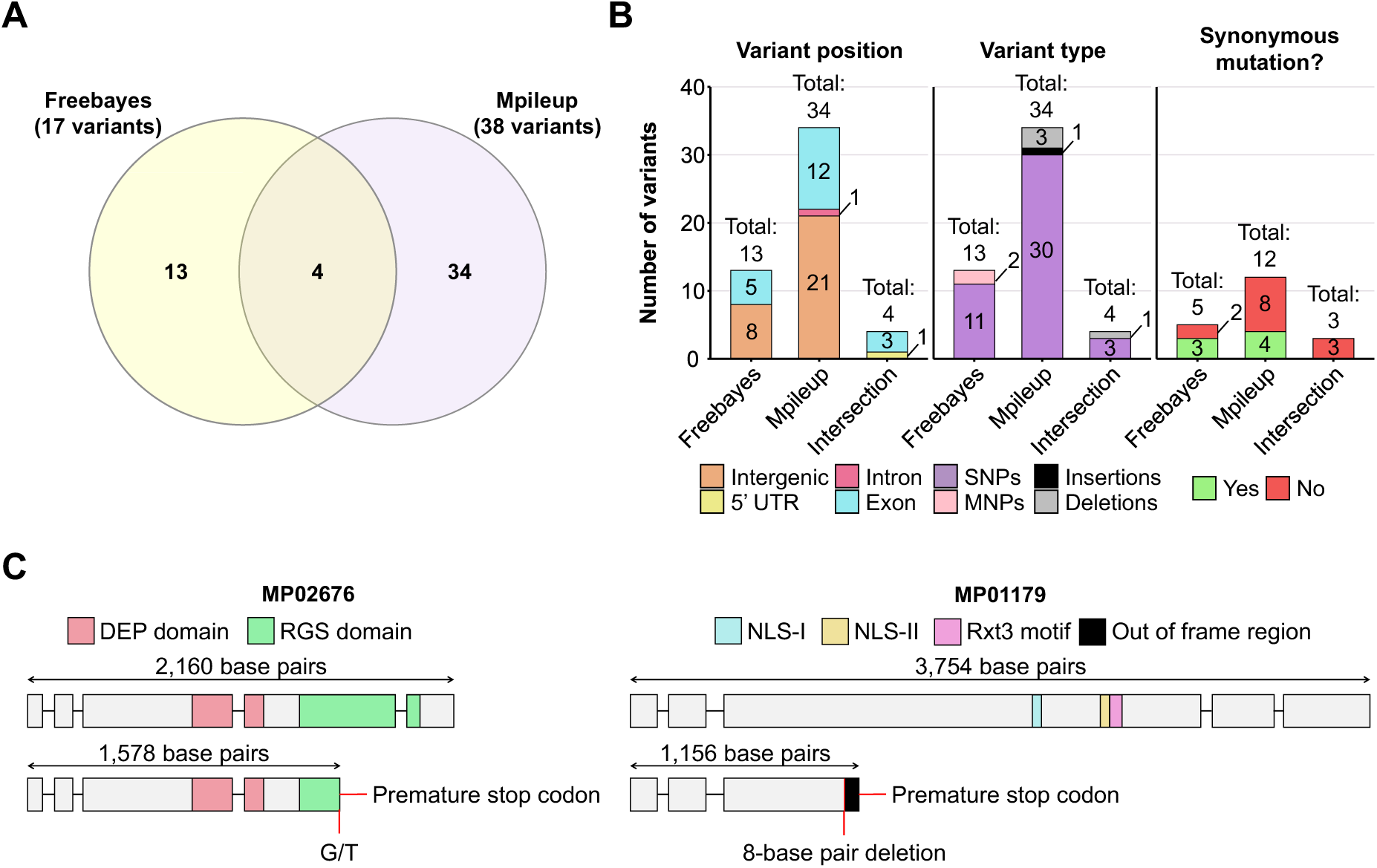
Identification of potential genomic variants associated with the FDS01 mutant phenotype. (A) Venn diagram showing the overlap of variants found by Freebayes and Mpileup when comparing FA553 and FDS01 genomes (B) Barplots indicating the number of variants uniquely identified by Freebayes, Mpileup and their intersection. The variants were classified by their position in *M. perniciosa* genome, their type and whether they encode an identical protein in both FA553 and FDS01 genomes (i.e., result in synonymous mutations; only applicable for SNPs in exons). (C) The MP02676 wild-type allele from FA553 and FDS02 isolates encodes a putative regulator of G protein signaling that possesses a DEP and an RGS domain. In the FDS01 genome, a SNP (G to T) results in a premature stop codon, producing a truncated protein with partial loss of the RGS domain, possibly affecting its function. The MP01179 wild-type allele encodes a putative transcriptional regulator containing two nuclear localization signals (NLS-I and NLS-II) and an Rxt3 motif. An 8-base pair deletion in the FDS01 genome results in a frameshift, leading to a premature stop-codon and a truncated protein. This truncation results in loss of both NLSs and the Rxt3 motif, likely compromising protein function. Exons are indicated by grey boxes, while introns are shown as lines. Supplementary Figure 6 shows these two mutations in reads derived from the FDS01 genome.

Among the three high-confidence SNPs identified in FDS01, one was located in the 5’ UTR of MP12249 (which encodes a putative terpene synthase) and another resulted in a non-synonymous mutation in MP10135. This mutation replaces a glutamate with a lysine in the encoded protein, which has no assigned function. The third SNP introduced a premature stop codon in MP02676 (Figure 8C), a gene that encodes a putative regulator of G-proteins. G-proteins control numerous cellular processes in fungi, including sexual and vegetative development (Wang et al. 2013; Park 2020). The early stop codon causes partial loss of the RGS (regulator of G-protein signaling) domain, which is required for G-protein inactivation (Wang et al. 2013; Park 2020).

Finally, the 8-nucleotide deletion in MP01179 (Figure 8C) also introduced a premature stop-codon, resulting in a truncated protein lacking 830 amino acids compared to its wild-type counterpart. Interestingly, this protein contains two predicted nuclear localization signals (NLSs) and a motif (Figure 8C) shared with the yeast Rxt3 protein, a component of the transcriptional repressor complex Rpd3 (Carrozza et al. 2005). Additionally, it shows 33.25% identity with a putative transcriptional repressor from *Coprinopsis cinerea* (GenBank: KAG2011931.1), and PSI-BLAST searches revealed 25% identity with the SAFB (Scaffold Attachment Factor Box)-Like transcriptional regulator from *Mus musculus*, which functions as a general transcriptional inhibitor (Chan et al. 2007). Thus, MP01179 likely encodes a novel transcriptional regulator that lost its function in the FDS01 mutant due to nucleotide deletions. This finding is consistent with the major transcriptional alterations observed in the FDS01 isolate (Figure 7).

## DISCUSSION

WBD is a major limiting factor for cacao cultivation in the Americas. Previous attempts to control the disease using strobilurin fungicides have been unsuccessful (Thomazella et al. 2012). Here, we show that *M. perniciosa* can tolerate high concentrations of strobilurins *in vitro*, and this tolerance is not explained by the typical mutations described in the *Cytb* gene (Supplementary Figure 3). Our results indicate that, in the presence of strobilurin, *M. perniciosa* undergoes intense metabolic reprogramming, including the repression of basic biological processes and activation of a series of catabolic processes and detoxification mechanisms.

Strobilurins inhibit complex III of the main respiratory chain, leading to ATP deprivation and oxidative stress, which can eventually cause fungal death (Balba 2007). The alternative oxidase enzyme, found in many phytopathogenic fungi (Tian et al. 2020), offers a bypass to complexes III and IV, alleviating the negative effects of strobilurin (Wood and Hollomon 2003). Indeed, the *M. perniciosa* AOX is important for strobilurin tolerance (Thomazella et al. 2012; Barsottini et al. 2019; Moretti-Almeida et al. 2019). Accordingly, we verified a rapid up-regulation of *Mp*-*Aox* upon azoxystrobin treatment (Figure 3A). In addition, an external NADH dehydrogenase gene (*NDH-2*) was induced by the fungicide (Figure 3A). Alternative non-proton pumping NADH dehydrogenases (e.g., NDH-2) are known to mitigate oxidative stress by promoting NADH oxidation, thus enhancing electron transport and metabolic flux (O’Donnell et al. 2011; Vamshi Krishna and Venkata Mohan 2019). While AOX and NDH-2 activity might promote fungal survival by reducing the generation of oxidative stress, ATP production is still negatively impacted due to their non-proton pumping nature (Vamshi Krishna and Venkata Mohan 2019; Tian et al. 2020). Importantly, these transcriptional alterations agree with the observed reduced growth of *M. perniciosa* mycelia in the presence of strobilurin (Figure 1).

To overcome azoxystrobin-induced ATP deprivation, *M. perniciosa* seems to activate important catabolic pathways. Genes encoding lipases and enzymes involved in both mitochondrial and peroxisomal ß-oxidation were strongly up-regulated in response to azoxystrobin, suggesting activation of lipid degradation (Supplementary Data Set 3). Interestingly, both pathways play important roles in the survival and pathogenicity of several fungi (Kretschmer et al. 2012; Patkar et al. 2012; Chen et al. 2016; Aliyu et al. 2019). NADH, FADH_2_ and acetyl-CoA are products of the ß-oxidation pathway. While NADH and FADH_2_ can be directly used in the electron transport chain to generate ATP, acetyl-CoA is oxidized in the TCA cycle to produce more reduced coenzymes and ATP (Houten and Wanders 2010). Consistent with this, genes coding for enzymes of the TCA cycle were also induced (Figure 4). Besides lipid catabolism, azoxystrobin appears to induce the degradation of branched-chain amino acids, which might generate additional acetyl-CoA for the TCA cycle, further contributing to ATP production (Supplementary Data Set 3) (Hildebrandt et al. 2015). These results suggest that *M. perniciosa* uses alternative energy sources to compensate for azoxystrobin-triggered ATP deprivation.

Genes encoding key enzymes of the glyoxylate cycle, isocitrate lyase (ICL) and malate synthase (MLS), were induced in *M. perniciosa* upon azoxystrobin treatment (Supplementary Data Set 3). These enzymes are exclusive to the glyoxylate cycle and generate intermediaries for the TCA cycle (Chew et al. 2019). Previous studies have described that these glyoxysomal enzymes play crucial roles on survival and pathogenicity of both plant and human pathogenic fungi. In *Candida albicans*, for example, the glyoxylate cycle enables yeast growth in nutrient-limited environments, within phagocytic cells (Dunn et al. 2009). Based on these findings, compounds with potential inhibitory effects on ICL have been proposed as drug targets for *C. albicans (Cheah et al. 2014)*. Likewise, azoxystrobin treatment triggers an energy-limiting condition in *M. perniciosa*, which results in the activation of the glyoxylate cycle. Therefore, our results suggest that the enzymes ICL and MLS might be prospective drug targets to be explored, potentially providing new options of drugs to be combined with azoxystrobin for the control of fungal diseases.

Even when exposed to high doses of azoxystrobin, *M. perniciosa* seems to produce enough energy to survive, albeit its growth rate is significantly reduced (Figure 1). This suggests that metabolic trade-offs may occur under fungicide exposure, favoring catabolism over anabolism for ATP generation. For instance, DNA, RNA and protein synthesis are primary consumers of ATP in a cell (Bennett et al. 2020). In *M. perniciosa*, genes involved in cell cycle progression, DNA replication, transcription and translation were down-regulated in the azoxystrobin-treated mycelia (Supplementary Data Set 4). Additionally, fungal genes required for ergosterol biosynthesis were repressed in response to the fungicide (Supplementary Data Set 4). Ergosterol is a major component of fungal cell membranes and is essential for maintaining membrane structure, fluidity and permeability (Rodrigues 2018). Therefore, inhibition of ergosterol biosynthesis negatively impacts fungal growth (Alcazar-Fuoli and Mellado 2013). For example, in *Fusarium graminearum*, the inhibition of ergosterol biosynthesis by thymol results in reduced growth rate (Gao et al. 2016). Similarly, azole antifungals target the 14α-demethylase enzyme (CYP51) in the ergosterol biosynthesis pathway, effectively inhibiting fungal growth (Ribas et al. 2016). In *M. perniciosa*, the possible repression of ergosterol biosynthesis by azoxystrobin (Supplementary Data Set 4) could compromise cell membrane integrity, contributing to the observed reduction in fungal growth.

*M. perniciosa* seems to activate a variety of detoxification mechanisms in response to azoxystrobin. Cytochrome P450 proteins are involved in a large diversity of biological processes, including degradation of xenobiotic compounds. Specifically, enzymes of the CYP53 subfamily are essential for the degradation of aromatic compounds, as they catalyze the para-hydroxylation of benzoate, a toxic intermediate (Jawallapersand et al. 2014). Our transcriptomic data showed that members of the CYP53 family are up-regulated by azoxystrobin (Figure 5C), which is an aromatic molecule. This suggests that the activation of these enzymes may be an attempt to detoxify the strobilurin fungicide. Notably, CYP53 family members are highly conserved in both ascomycetes and basidiomycetes and are critical for *Aspergillus nidulans* survival in the presence of benzoate (Fraser et al. 2002). Furthermore, no homologs of this family is present in higher eucaryotes (Jawallapersand et al. 2014), making these enzymes potential drug targets. Drug transporters may also contribute to *M. perniciosa* detoxification response to azoxystrobin. In fungi, transporters can function as efflux pumps, expelling xenobiotic compounds into the external environment, thereby preventing their toxic accumulation inside the cells and contributing to pathogen insensitivity or resistance to fungicides (de Waard et al. 2006). The major superfamilies of drug transporters, 2.A.1 (MFS) and 3.A.1 (ABC), are often associated with multi-drug resistance in phytopathogenic fungi (Hu and Chen 2021). Genes encoding proteins from both superfamilies were activated in the *M. perniciosa* mycelia treated with azoxystrobin (Figure 5B). Additionally, a total of 15 genes encoding glutathione S-transferases (GSTs) were up-regulated in response to azoxystrobin (Figure 5C). These enzymes are involved in the detoxification of xenobiotic compounds and protection against oxidative stress (Hayes et al. 2005). Moreover, GSTs have been linked to resistance against the fungicides chlorothalonil and benzimidazole (Shin et al. 2003; Sevastos et al. 2017). Therefore, our results suggest that *M. perniciosa* may also employ GSTs to mitigate the cytotoxic effects of azoxystrobin (Figure 5C).

The indiscriminate use of fungicides can drive the emergence of resistant fungal strains. The lack of rotation in fungicide application and prolonged use of chemicals have contributed to the emergence of resistant isolates in several fungal pathogens (Lucas et al., 2015). Antimicrobials often expose pathogens to DNA damage and mutagenesis (Shapiro 2015; Amaradasa and Everhart 2016; Gambhir et al. 2021), although these effects can be usually mitigated by DNA repair systems (Chatterjee and Walker 2017). However, the deregulation of these protective systems can lead to increased genomic instability and the emergence of drug resistance (Legrand Melanie et al. 2007; Boyce Kylie J. et al. 2017). Interestingly, azoxystrobin treatment down-regulated genes related to DNA repair (Supplementary Data Set 4), which may lead to genetic diversification and the emergence of fungicide resistance. Indeed, after long-term exposure of *M. perniciosa* to azoxystrobin under laboratory conditions, we isolated a mycelial sector displaying vigorous growth (named FDS01), suggesting the spontaneous emergence of a fully resistant mutant (Figure 6). Unlike the parental strain, which tolerates strobilurins but shows reduced growth rate, growth of the FDS01 mutant is not affected by azoxystrobin. Although single-point mutations in the *CytB* gene are typically responsible for resistance to strobilurin fungicides (Fernández-Ortuño et al. 2008), no mutations were found in the *CytB* gene of FDS01 (Supplementary Figure 3). Furthermore, AOX activity, known to enhance strobilurin tolerance (Thomazella et al. 2012), did not differ in expression between the resistant mutant and wild-type isolates FA553 and FDS02 (Supplementary Figure 5). This suggests that the FDS01 resistance to strobilurins may be due to a novel, non-canonical mechanism.

Genome analysis of FDS01 revealed an 8-nucleotide deletion in a putative transcription factor (MP01179 gene), as well as a non-synonymous mutation in an RGS-like protein (MP02676 gene), both of which could explain the mutant phenotype (Figure 8). The deletion in the gene MP01179 produces a premature stop codon resulting in a truncated protein with only 335 of the 1165 amino acids present in the wild-type protein. This truncated version lacks both the two predicted NLSs (nuclear localization signals) and the putative Rxt3 motif (Figure 8). The absence of NLSs would prevent the protein from entering the nucleus, thus abolishing its function as a transcriptional regulator (Lu et al. 2021). Deletion of the Rxt3 subunit from the Rpd3L histone deacetylase complex of *Aspergillus oryzae* results in enhanced tolerance to multiple stresses, including osmotic and oxidative stress (Chang et al. 2024). Thus, loss of the Rxt3 motif from MP01179 may contribute to the altered transcriptional program of the FDS01 isolate and its enhanced azoxystrobin resistance, further pointing to the pivotal role of transcriptional regulation in the development of fungicide resistance (Thakur et al. 2008; Liu et al. 2015; Hu et al. 2018; Sang et al. 2018)

The MP02676 gene encodes a protein containing DEP and RGS domains, which are typical of repressors of G-protein signaling (Segers Gerrit C. et al. 2004; Wang Ping et al. 2004), a class of molecules reported to have distinct roles in fungi (Wang et al. 2013; Park 2020). These repressors use their RGS domain to deactivate G-proteins (Wang et al. 2013; Park 2020). Interestingly, the SNP found in MP02676 produces an early stop codon, leading to the partial loss of the RGS domain (Figure 8C), which may compromise its function. In *Aspergillus nidulans* and *A. fumigatus*, the FlbA RGS has a major role in growth regulation, and its deletion results in increased hyphal growth (Tamame M et al. 1983; Yu et al. 1996; Mah and Yu 2006). Although MP02676 shares only 34.55% similarity with *A. nidulans* FlbA (Uniprot: P38093.1), the partial loss of the MP02676 RGS domain in FDS01 could similarly contribute to its vigorous hyphal growth in the presence of azoxystrobin (Figure 6).

The hypotheses regarding the contribution of the mutated versions of MP01179 and MP02676 to the mutant phenotype could be tested through genetic complementation of FDS01 with the wild-type allele or by generating independent mutants of each of these genes. However, an efficient genetic manipulation technique for *M. perniciosa* has not yet been developed. Thus, despite the available evidence, we are currently unable to confirm that the loss of function of MP01179 and MP02676 confers resistance to azoxystrobin. Yet, the results and functional evidence we provide here may guide future studies related to these genes in either *M. perniciosa* or other fungal species. Importantly, future advancements in genetic manipulation of *M. perniciosa* will allow for the confirmation of our hypotheses.

In conclusion, our results suggest that *M. perniciosa* shifts its metabolism towards an energy-saving state and activates distinct detoxification processes to cope with azoxystrobin toxicity. Although this comes with a reduced growth rate, it enables fungal survival and may favor the emergence of resistant strains. This study underscores the importance of combining molecules with different modes of action for effective pathogen control. In addition, designing hybrid bifunctional compounds that target independent molecules or pathways in the pathogen might also offer a promising strategy to improve fungicide efficiency and durability in agriculture (Zuccolo et al. 2019). Future studies will investigate the contribution of azoxystrobin-responsive genes to *M. perniciosa* tolerance to strobilurins, aiming to confirm if they are viable targets for developing new fungicides.

## ACKNOWLEDGEMENTS

This study was supported by the São Paulo Research Foundation (FAPESP) through fellowships awarded to P.F.V.P, C.V.C.M., G.L.F. and D.P.T.T. (process numbers 20/04773-8, 14/00802-2, 14/06181-0, 13/05979-5 and 12/09136-0). B.A.P. was financed by the Coordenação de Aperfeiçoamento de Pessoal de Nível Superior – Brasil (CAPES) – Finance Code 001. P.J.P.L.T received funds from the National Council for Scientific and Technological Development (CNPq; grants 308349/2022-9 and 443671/2018-4), the Serrapilheira Institute (grant G-1811- 25705), and FAPESP (grant 18/24432-0). We thank the University of North Carolina High Throughput Sequencing Facility for their support in providing sequencing services and training, as well as Natália Mincov Costa for her technical support and assistance in maintaining *M. perniciosa* cultures.

## Supplementary Figures

**Supplementary Figure 1.**
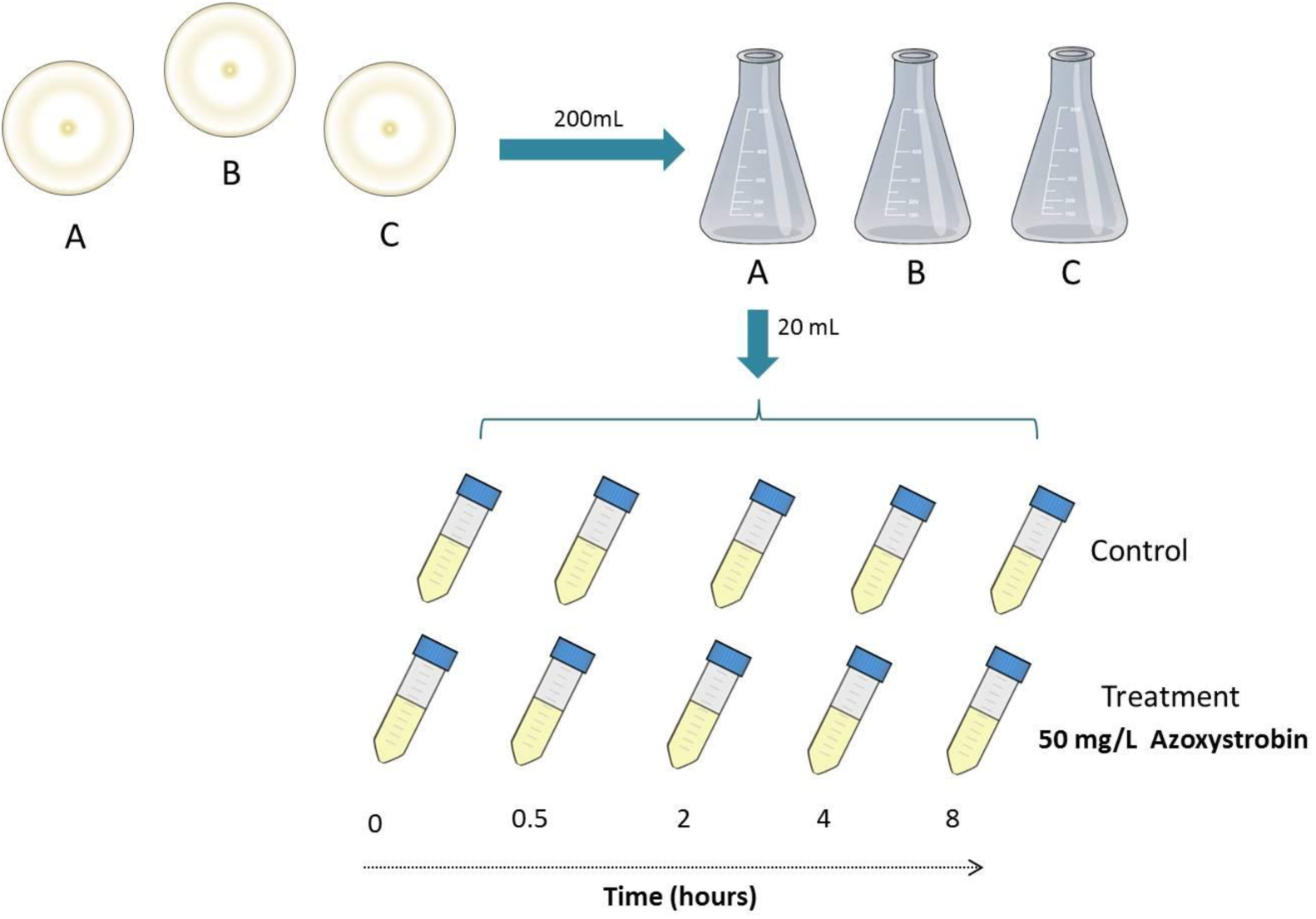
Experimental design for *M. perniciosa* treatment with azoxystrobin in a time-course experiment. Three biological replicates were inoculated in liquid Malt light medium. Samples for RNA sequencing were collected at five time points after azoxystrobin treatment, along with corresponding control samples (without fungicide). The experiment comprises a total of 30 individual RNA-seq libraries per isolate.

**Supplementary Figure 2.**
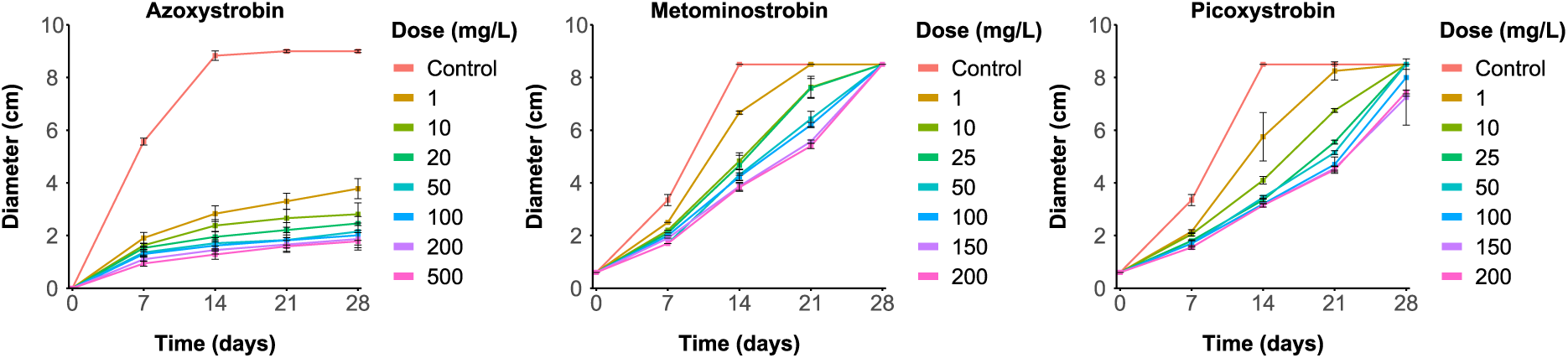
Evaluation of *M. perniciosa* growth in the presence of the strobiluins azoxystrobin, metominostrobin and picoxystrobin. The diameter of fungal colonies was measured under different concentrations of azoxystrobin (left), metominostrobin (center) and picoxytrobin (right) over 28 days. Error bars represent the standard error (n= 3). In the control conditions (i.e., without fungicide), the mycelium covered the plate (9 cm in diameter) in approximately 14 days, leading to the plateau observed in the graphs.

**Supplementary Figure 3.**
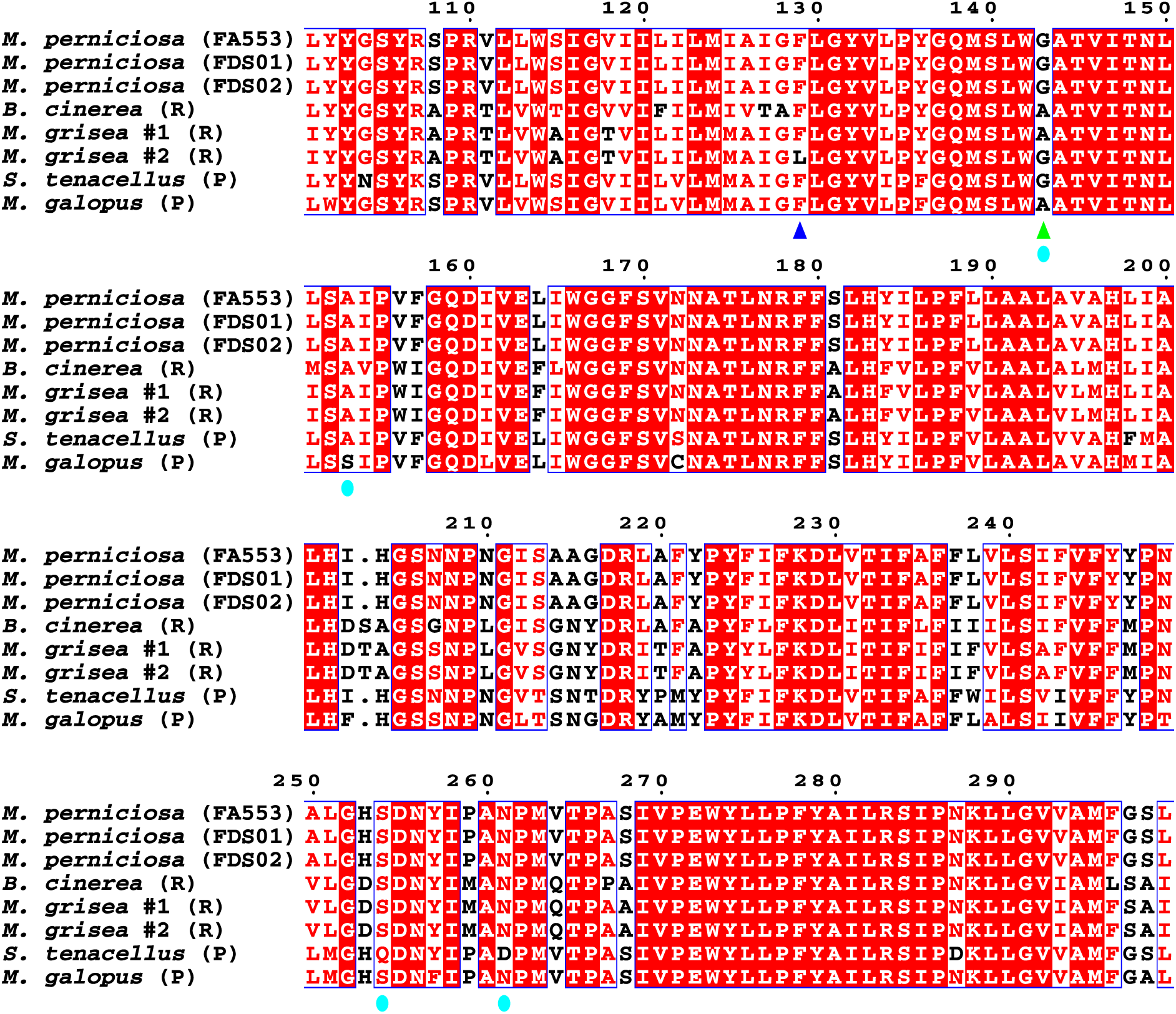
Mutations in the *CytB* gene that determine azoxystrobin resistance in phytopathogens are absent in *M. perniciosa*. Alignment of a 300-amino acid fragment of the CytB (cytochrome B) protein from different fungi. Dark blue and green arrows correspond to the F129L and G143A mutations, respectively, which are known to confer strobilurin resistance in pythopathogens (R). Cyan circles indicate mutations in the CytB protein that have been identified in strobilurin-producing basidiomycetes (P) and confer natural resistance to their own product. None of the *M. perniciosa* isolates used in this study contain any of these alterations. Protein sequences were retrieved from the NCBI database with the following IDs: *Botrytis cinerea* (ACL50592.1); *Magnaporthe grisea* G143A mutant (AAO91631.1); *Magnaporthe grisea* F129L mutant (AAO91630.1); *Strobilurus tenacellus* (CAA61250.1); *Mycena galopus* (CAA61247.1).

**Supplementary Figure 4.**
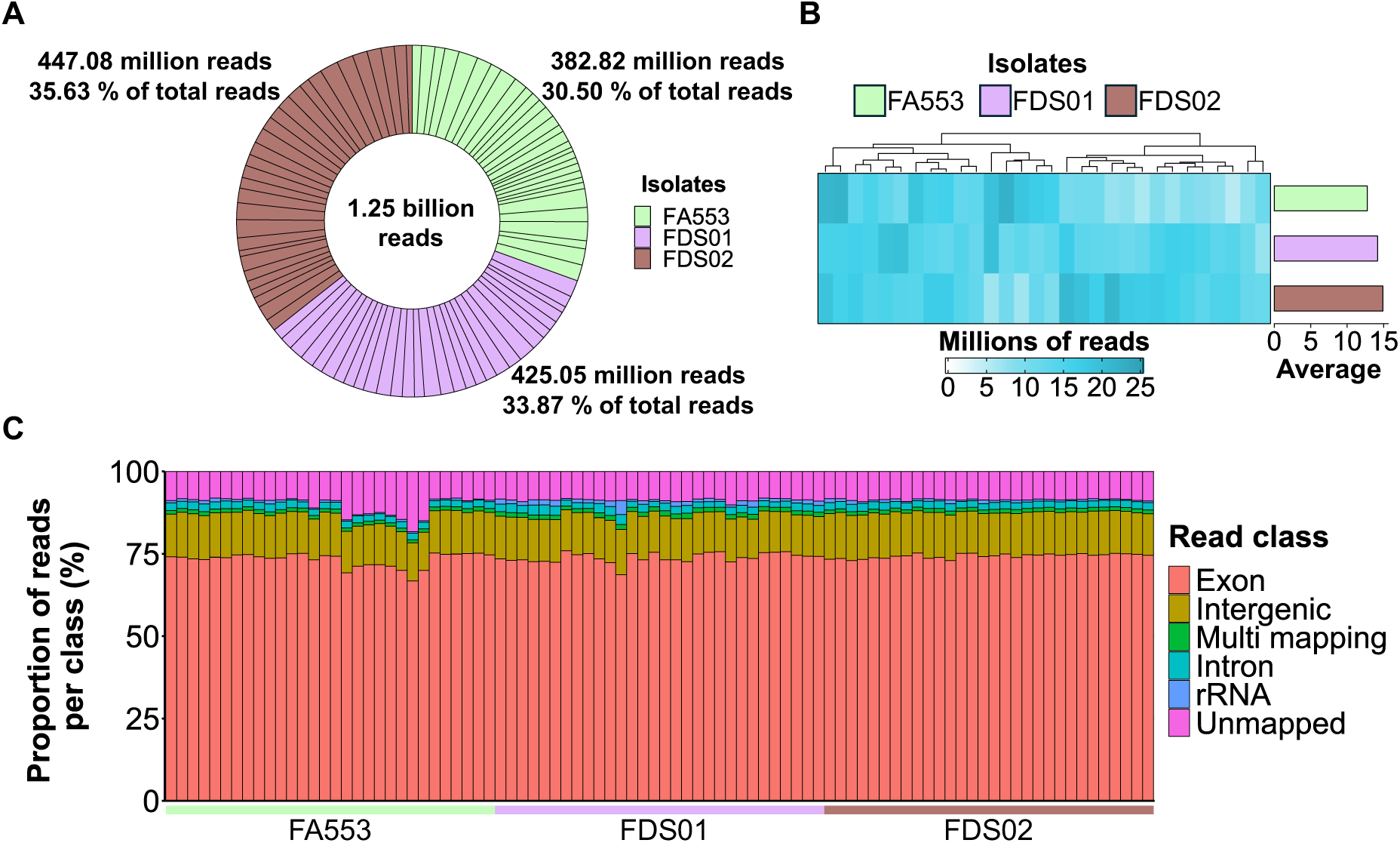
Sequencing and mapping metrics. (A) Distribution of the 1.25 billion reads generated for the 90 RNA-seq libraries from three *M. perniciosa* isolates (FA553, FDS01 and FDS02). Each sector in the chart represents an individual library. (B) Heatmap showing the number of reads per RNA-seq library. Each row corresponds to one *M. perniciosa* isolate, and the bar plots on the right show the average number of reads produced for each library. (C) Percentage of reads mapped to intergenic and genic regions (exon and intron), multi-mapped to more than one genomic position, mapped to rRNA and unmapped reads (total reads passing QC minus mapped reads). Each library is represented by a column.

**Supplementary Figure 5.**
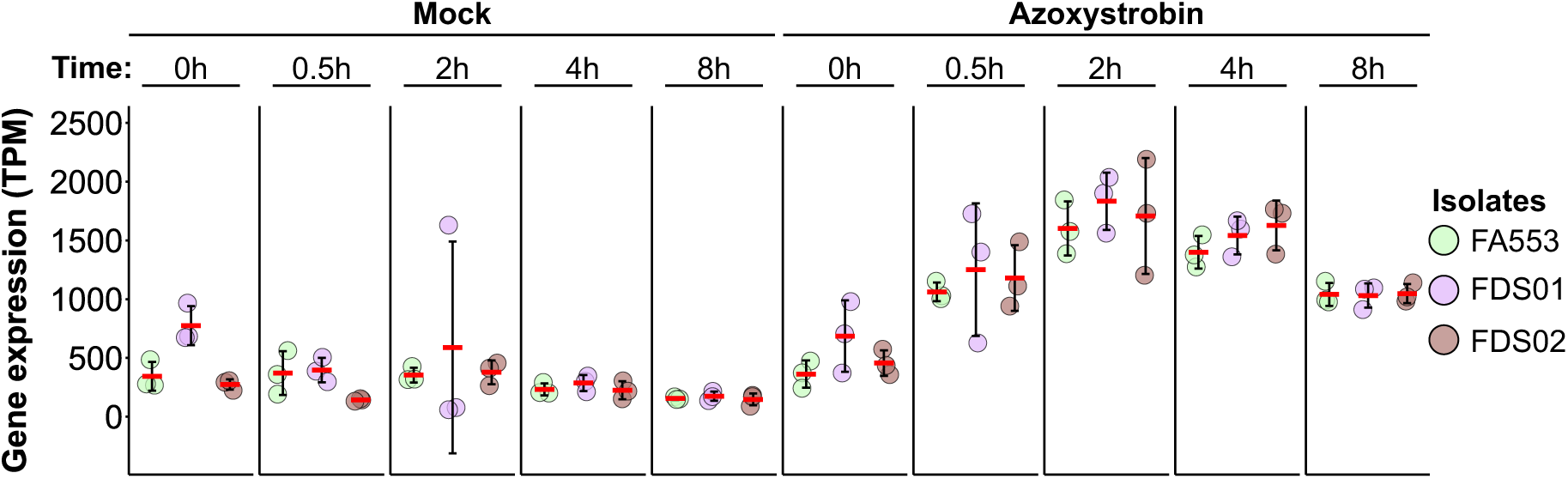
Up-regulation of alternative oxidase (*Mp*-*Aox*) is not linked to the increased tolerance of the FDS01 isolate to azoxystrobin. *Mp*-*Aox* is similarly up-regulated in all three *M. perniciosa* isolates at all time points.

**Supplementary Figure 6.**
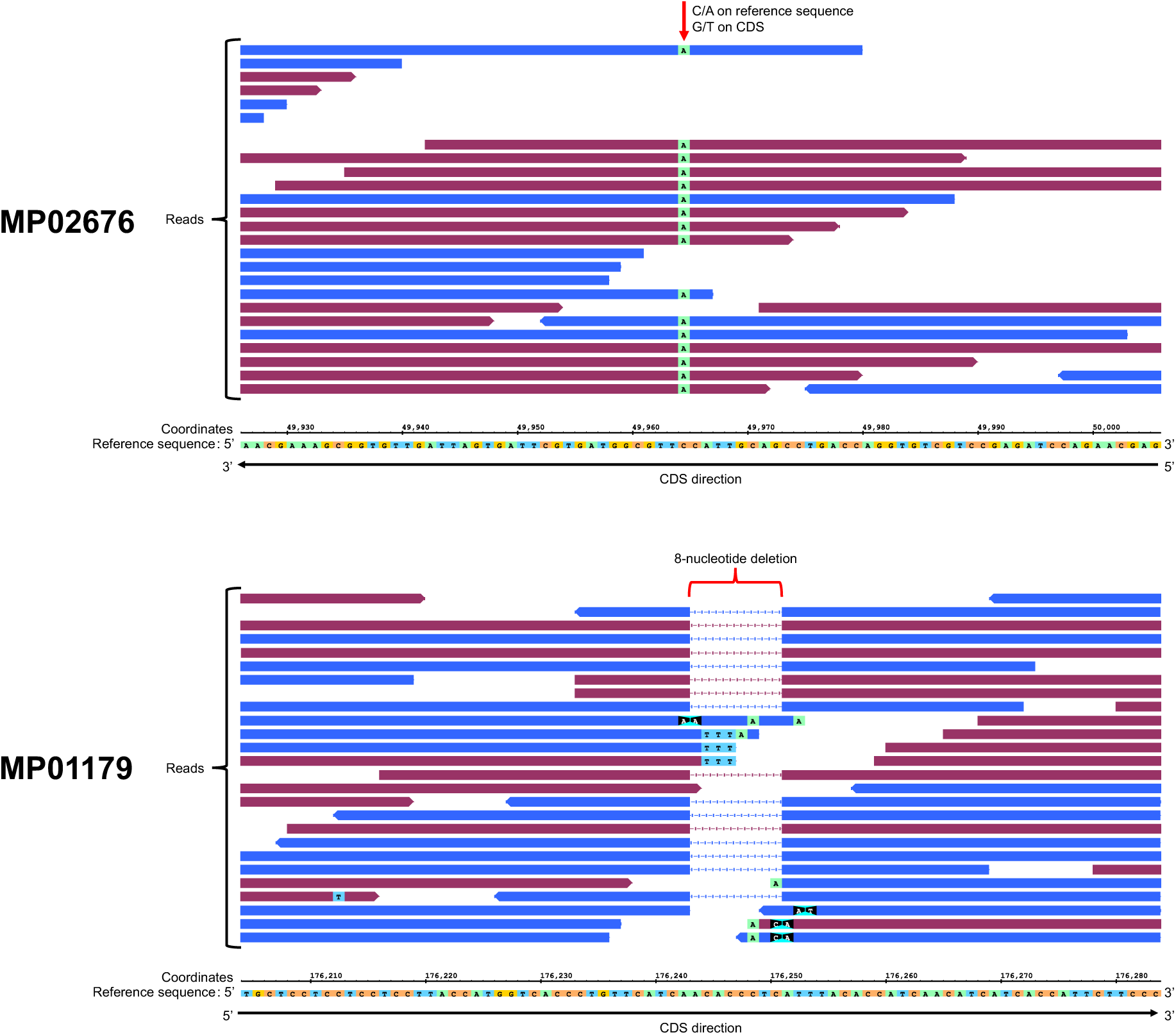
Genome sequencing reveals mutations in two genes of the FDS01 isolate. Sequenced reads of the MP02676 gene from the FDS01 genome show the occurrence of a SNP (C to A in the reference sequence and G to T in the reference CDS) compared to the FA553 reference genome. Additionally, gaps in the FDS01 genomic reads indicate an 8-nucleotide deletion in the MP01179 gene.

## Notes

### Competing Interest Statement

The authors have declared no competing interest.

